# Extending sampling approaches for great crested newt (*Triturus cristatus*) eDNA monitoring

**DOI:** 10.1101/2024.10.21.619420

**Authors:** Lynsey R. Harper, Kirsten J. Harper, Suzie Platts, Rebecca Irwin, Michael Bennett, Ben Jones, Oliver Taylor, Laura Plant, Danielle Eccleshall, Andrew Briscoe, Luke Gorman, Kat Stanhope, Bastian Egeter

## Abstract

1. Environmental DNA (eDNA) monitoring has been used for great crested newt (*Triturus cristatus*) survey in the UK since the publication of a Defra-funded trial in 2014. If eDNA results are to be used in support of a great crested newt licence, surveys must be performed during a 76-day survey window (15 April – 30 June) to coincide with peak great crested newt activity, and must follow the approved ethanol precipitation protocol. However, eDNA detection is possible in other months and filtration may be a more effective method of eDNA capture.
2. We investigated whether the great crested newt eDNA survey season could be extended and filtration could be used for great crested newt eDNA capture by reviewing the available evidence and conducting a field study from April to October 2022. Paired water samples for ethanol precipitation and filtration were collected from 25 ponds once a month, resulting in 124 samples of each type. All samples (N = 248) were analysed with the approved great crested newt quantitative PCR assay.
3. Our results indicate that great crested newts can be reliably detected using both eDNA capture methods from April to August, with detection rates decreasing in September and October. Great crested newt eDNA detection was comparable or higher with filtration than ethanol precipitation.
4. *Practical implication.* Acceptance of filtration for great crested newt eDNA surveys could allow more water to be processed for robust and reliable estimates of great crested newt presence. Extending the great crested newt eDNA survey season to August could allow more waterbodies to be surveyed for great crested newt presence (but not absence), and identification of sites that provide important habitat for great crested newts outside of the breeding season. This would also remove logistical challenges and costs associated with completing sampling within 11 weeks and laboratory analysis within 10 working days from sample receipt. Furthermore, great crested newt eDNA surveys could be more frequently carried out alongside monitoring for other species, which are typically surveyed from April to September/October or year-round with conventional methods or eDNA surveys using filtration. This could enable infrastructure projects to develop more effective mitigation measures as well as reduce time required from surveyors and survey costs.

## 1. Introduction

Great crested newts (*Triturus cristatus*) are a European protected species, meaning individuals, their eggs, their breeding sites, and their resting places are protected by law. In the UK, licences must be obtained for activities affecting great crested newts, including development and mitigation, from the appropriate statutory authority, e.g. Natural England, NatureScot or Natural Resources Wales. Where records suggest great crested newts may be present, there is suitable aquatic habitat, or suitable terrestrial habitat at or in the vicinity of a proposed development site, surveys are required to assess species distribution and abundance in order to take appropriate measures to avoid, mitigate or compensate for any negative effects on great crested newts. Traditionally, great crested newts were monitored using methods such as torchlight counts, bottle trapping, netting and egg searches to confirm presence or likely absence in waterbodies (Langton et al., 2001). Since 2014, environmental DNA (eDNA) analysis has offered a non-invasive, time-efficient and cost-effective alternative (Biggs et al., 2015; Rees et al., 2014), but within a regulated 76-day survey window (15 April – 30 June) to coincide with peak breeding activity and maximise chance of detection (Biggs et al., 2014).

Previous UK studies have shown that great crested newts can be successfully detected with eDNA analysis outside this regulated survey window (Buxton et al., 2017, 2018a; Gorman et al., 2020; Rees et al., 2017), possibly due to juveniles remaining beyond the breeding season, and juveniles and adults overwintering in waterbodies (Rees et al., 2023). Across Europe where the same regulations do not apply, eDNA surveys for great crested newts have also occurred over summer months (Knudsen et al., 2023; Strand et al., 2022). Extension of the eDNA survey window in the UK could allow more potential great crested newt waterbodies to be surveyed for robust and reliable estimates of great crested newt presence. It could also allow great crested newt eDNA surveys to be more frequently carried out alongside monitoring for other species, which are typically surveyed from April to September/October or year-round with conventional methods or eDNA surveys using filtration. Potential pairings may include smooth newt (*Lissotriton vulgaris*), chytrid fungus (*Batrachochytrium dendrobaditis*; Taugbøl et al., 2021, 2025), and white-clawed crayfish (*Austropotamobius pallipes*; Troth et al., 2020), or amphibian assemblages (Charvoz et al., 2021; Davison et al., 2025; Dufresnes et al., 2019; Strand et al., 2022; Svenningsen et al., 2022) or whole vertebrate communities (L. R. Harper et al., 2018). This could enable infrastructure projects to develop more effective mitigation measures as well as reduce survey time and costs. However, additional evidence is needed to support extension of the great crested newt eDNA survey window and regulatory change (Rees et al., 2017, 2023).

Technical guidance published by Biggs et al. (2014) and approved by Natural England underpins the widely adopted protocol (WC1067) for great crested newt eDNA surveys in the UK. This guidance was based on a project conducted in 2013/14 that established the performance of and produced a technical advice note on eDNA techniques for the great crested newt. In accordance with this guidance, all great crested newt eDNA sampling kits produced by different commercial eDNA providers capture eDNA via ethanol precipitation (hereafter EP). This method purifies and concentrates DNA by adding salt and ethanol to water samples, which forces the precipitation of DNA out of the solution. The DNA can then be separated from the rest of the solution by centrifugation. EP is restricted to small water volumes (e.g. 90 mL) and susceptible to contamination during laboratory processing. It can also be expensive due to the requirement for molecular biology grade ethanol, which: can be difficult to procure; can be subject to taxes (unless procured under a duty-free licence); is classed as dangerous goods for transportation necessitating specialist couriers, specific packaging and limited quantities for shipping (especially costly for small numbers of kits); and must be stored appropriately with specialist waste disposal. EP kits potentially present a risk to ecologists and any others who have to store and transport a large number of kits (Bruce et al., 2021; Rees et al., 2023).

Since 2014, eDNA applications have been continually evolving and multiple studies have identified that filtration may provide more efficient species detection than EP (e.g. Hinlo et al., 2017; Minamoto et al., 2015; Peixoto et al., 2020; Spens et al., 2017; Troth et al., 2020), although the optimal capture method can depend on environment sampled and inhibitor concentrations (Minamoto et al., 2015; Troth et al., 2020). Filtration separates DNA from water using a filter medium (e.g. paper, gauze) that allows the solution to pass through but not the DNA. Filters can possibly process litres of water for higher detection rates, albeit less water may be processed and/or processing may take longer for turbid water samples (Andreou et al., 2023; Minamoto et al., 2015). Unlike EP samples, filters are self-contained and quicker to extract for lower contamination risk and reduced labour costs. There is minimal risk to ecologists and any others who have to store and transport them, meaning standard couriers can be used for shipping. They can also be stored at ambient temperature before and after (if a preservative solution is added to the filter) use, making storage of collected samples easier and less expensive (Bruce et al., 2021; Rees et al., 2023). In contrast, EP kits must be stored at 2-4°C if they cannot be delivered to a laboratory on the same day as sample collection (Biggs et al., 2014).

Here, we reviewed the evidence for extending the great crested newt eDNA survey season and using filtration for great crested newt eDNA capture. We performed a field comparison of in-season (April to June) versus out-of-season (July to October) eDNA survey, and EP versus filtration for great crested newt eDNA detection. We hypothesised that great crested newts would be detected beyond 30 June, and that filtration would outperform EP for great crested newt eDNA detection. We provide evidence-based recommendations for regulatory changes to evolve and streamline great crested newt eDNA surveys.

## 2. Materials and Methods

### 2.1. Literature review

A literature review was undertaken in June 2022 to synthesise published and grey literature on using filtration for great crested newt eDNA capture and great crested newt eDNA detection out-of-season. Published literature was searched with Google Scholar using the search terms “environmental DNA”, “eDNA”, “ethanol precipitation”, “filtration”, “qPCR”, “great crested newt”, “*Triturus cristatus*”, “GCN”, and “amphibians”. Literature was restricted to those referencing great crested newts as well as incorporating filtration and/or out-of-season eDNA surveys. Grey literature was sought by contacting a range of regulatory agencies, non-governmental organisations, and ecological consultancies. Nineteen organisations were contacted, including five regulatory agencies, three non-governmental organisations, and 11 ecological consultancies. Of these, 12 responded, including three which had relevant data but only two provided. Cited literature within studies returned by searches, including previous literature reviews and reports, was also accessed and assessed for relevance. Information extracted from each source was summarised in Microsoft Excel (Supporting Information: Appendix 1).

### 2.2. Field study

Ponds across three sites in Buckinghamshire, England, were selected based on results of previous surveys conducted on behalf of HS2 Ltd within the last four years but dating back to 2013 for some ponds (Table 1, Figure 1). Surveys had not been undertaken for two ponds included in the present study. Most ponds were created as part of ecological mitigation works and were receptor sites for great crested newt translocation. Ponds were located near Newton Purcell (Ponds 1-4), Great Missenden (Ponds 5-8), and Aylesbury (Ponds 9-25). The Newton Purcell and Great Missenden sites were over 35 km apart, with the Aylesbury site in between (over 20 km from Newton Purcell site and over 15 km from Great Missenden site). Therefore, great crested newt dispersal between the three sites is highly unlikely. However, most ponds within each site were less than 500 m apart and within similar proximity to other ponds not included in this study, therefore great crested newts at each site could move between ponds in response to environmental and social cues (Unglaub, Cayuela et al., 2021).

**Figure 1.**
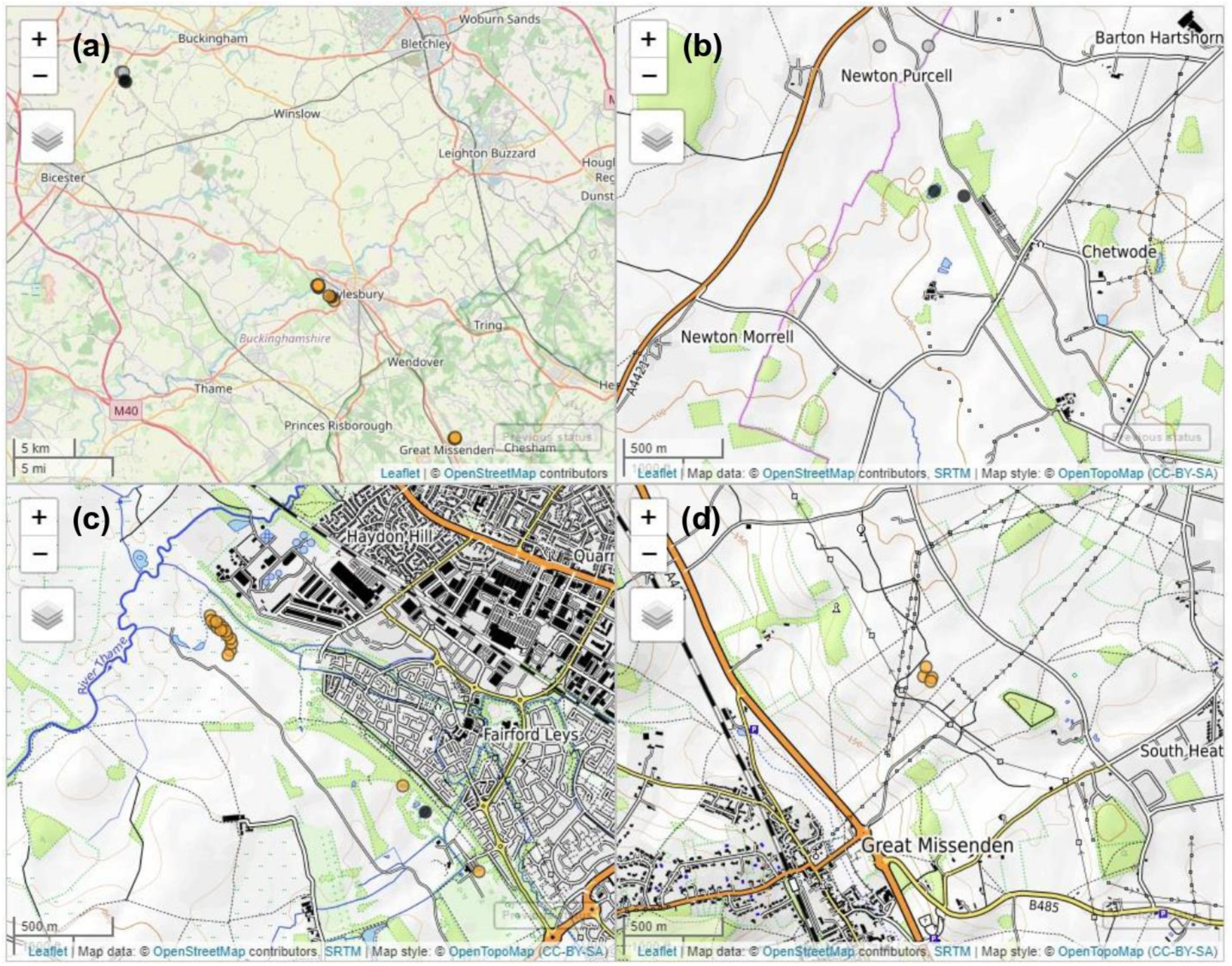
Maps of study ponds in Buckinghamshire. All ponds are shown relative to the wider landscape **(a)**, then by clusters near Newton Purcell **(b)**, Aylesbury **(c)**, and Great Missenden **(d)**. Points are coloured by previous great crested newt status, where orange is positive, black is negative, and grey is unknown.

**Table 1.**
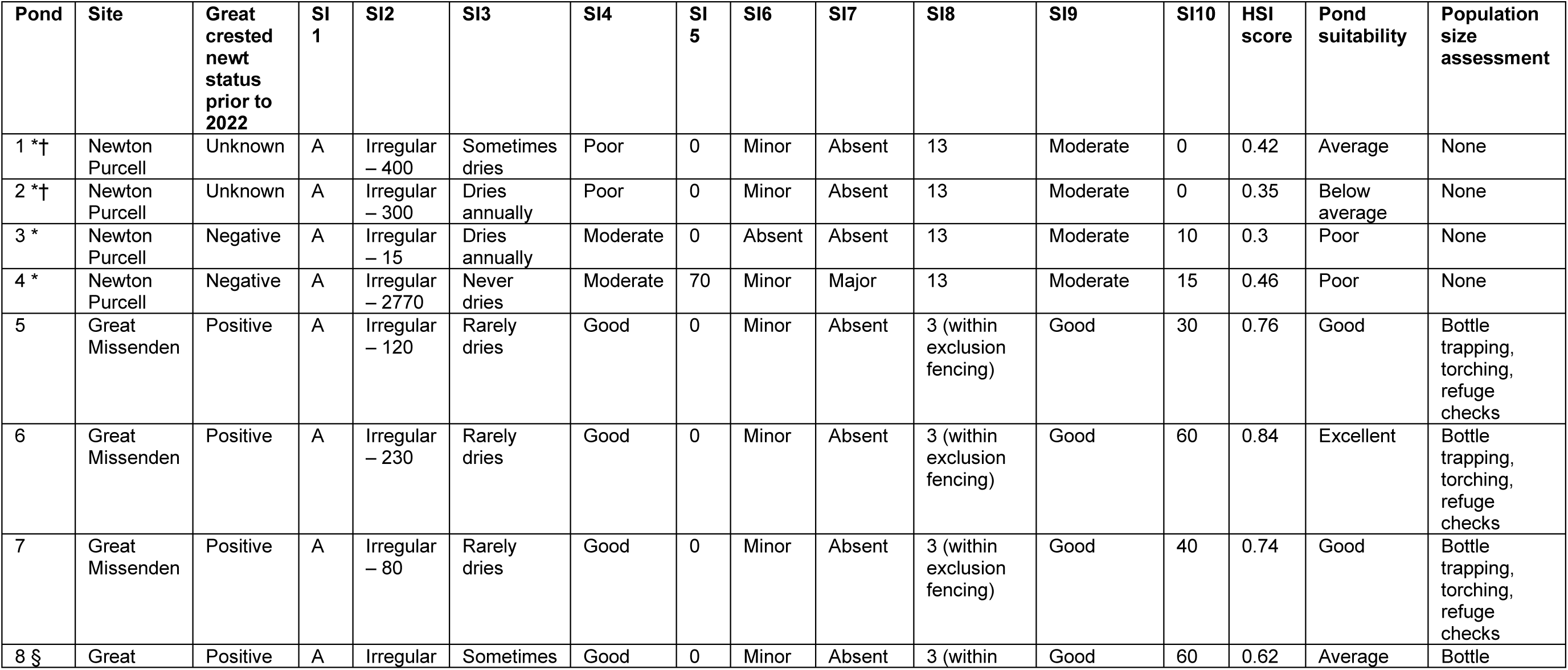
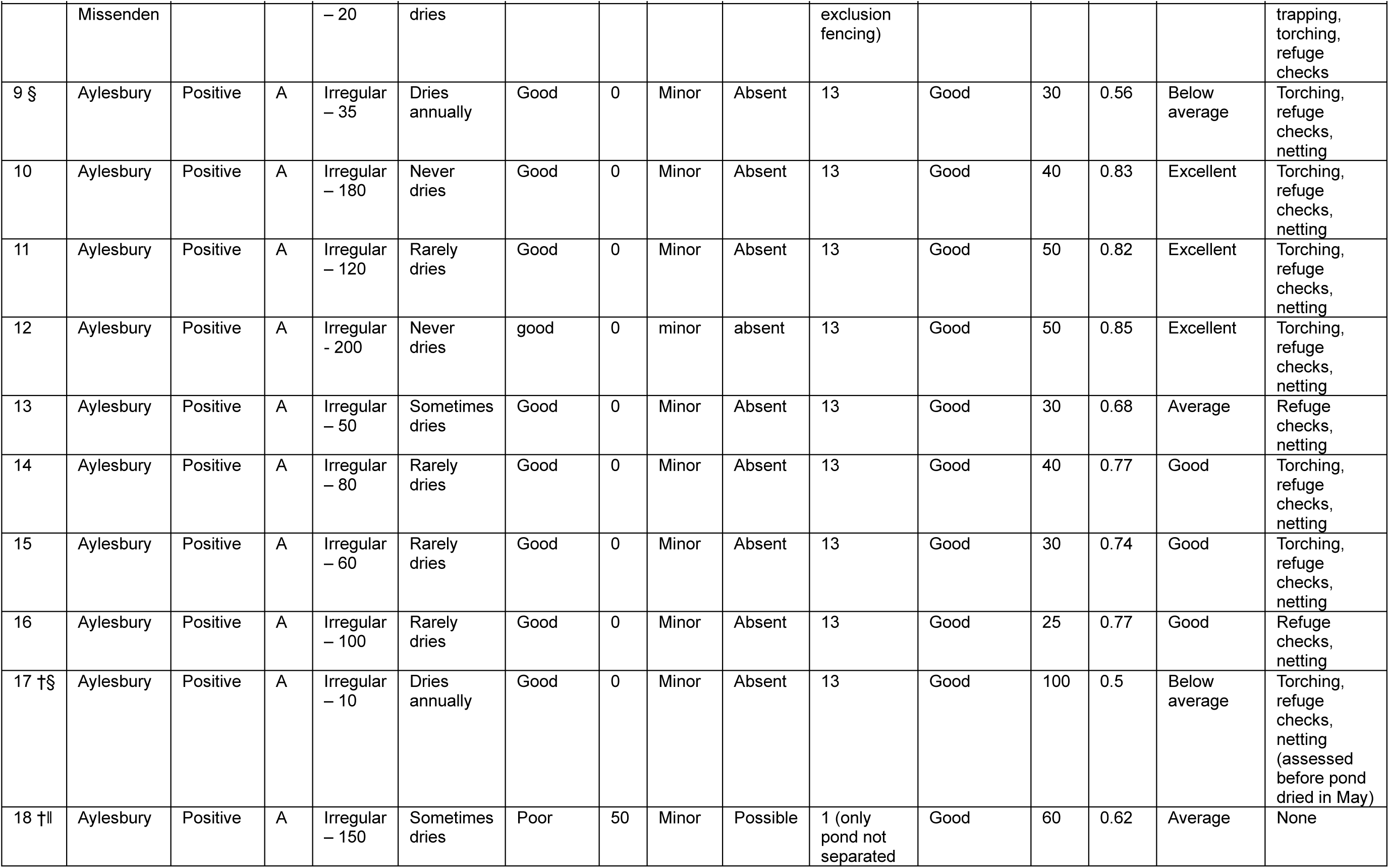
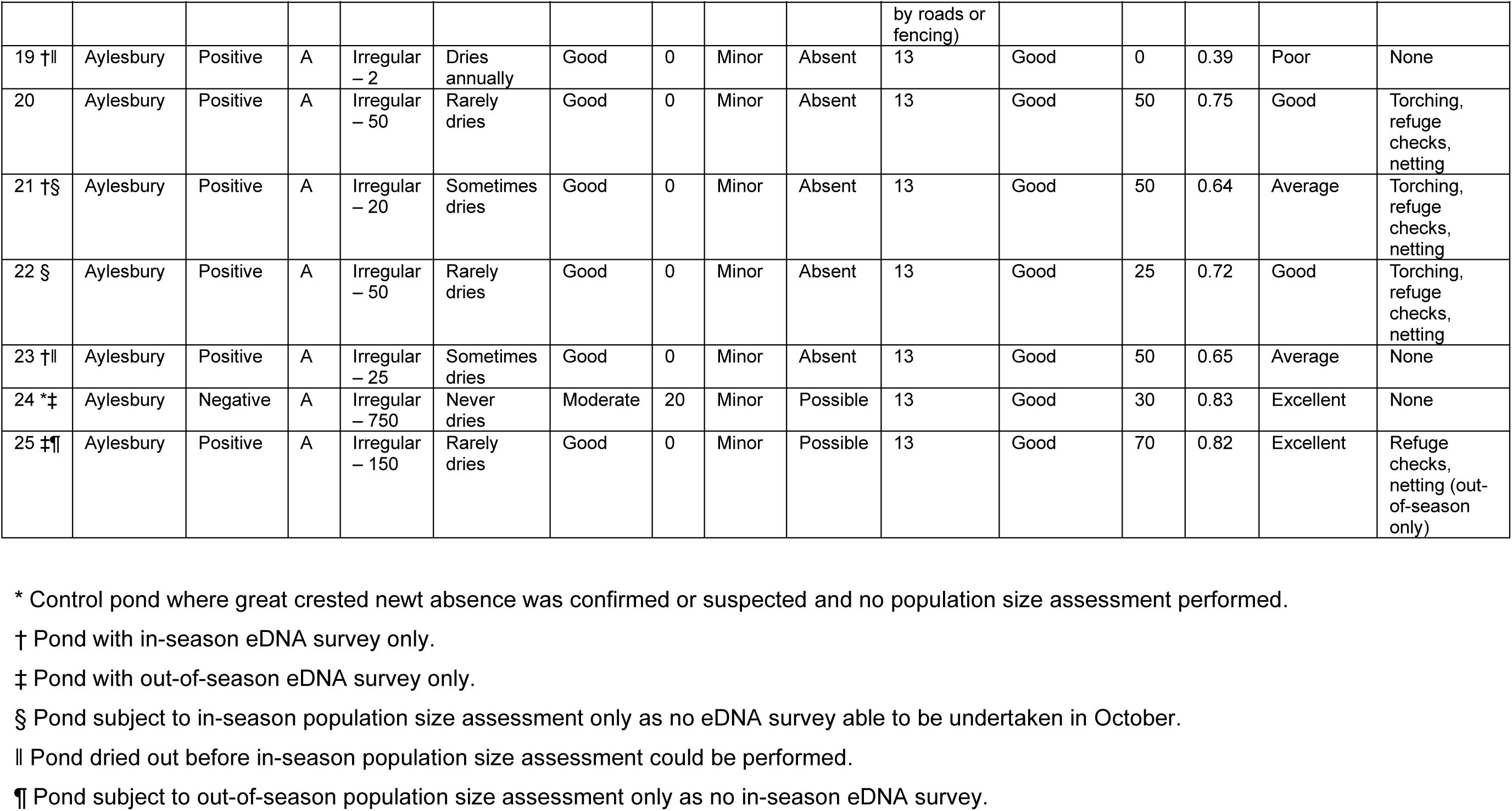
Characteristics of study ponds based on the ten suitability indices (SI) that are used to calculate the Habitat Suitability Index (HSI) for great crested newts (Oldham et al., 2000), and define pond suitability (ARG-UK, 2010). SI1: Geographic location (Zone A = optimal, Zone B = marginal, Zone C = unsuitable). SI2: Pond area (rectangle/ellipse/irregular – m^2^). SI3: Permanence (never dries/rarely dries/sometimes dries/dries annually). SI4: Water quality (good/moderate/poor/bad). SI5: Shade (percentage of margin shaded 1 m from bank). SI6: Waterfowl (absent/minor/major). SI7: Fish (absent/possible/minor/major). SI8: Pond count (number of ponds within 1 km). SI9: Terrestrial habitat (good/moderate/poor/none). SI10: Macrophytes (percentage of the pond surface area occupied by macrophyte cover).

Initially, 21 ponds (ponds 1-2, 4-17 and 19-23) were sampled in April 2022 to ensure at least 17 positive ponds (ponds 5-17 and 19-23) and three negative ponds (i.e. control ponds, ponds 1, 2 and 4) for great crested newts. From May onwards, 20 ponds (ponds 1, 3-16, 18, and 20-23) were selected for repeat surveys (ponds 2, 17 and 19 had dried). Over the study period, ponds from the same three sites were added (ponds 3, 18, 24 and 25) to replace any that dried and/or became unsafe to access between April and June (ponds 1, 2, 17, 19 and 23). This resulted in a total of 25 ponds being surveyed over the study period, but ultimately the number of ponds sampled each month decreased (Supporting Information: Appendix 2).

Each pond was sampled by AtkinsRéalis ecologists once each month using both NatureMetrics Great Crested Newt eDNA Kits (EP) and Pump Aquatic eDNA Kits (filtration; Figure 2). The latter included a sterile 0.8 μm pore size polyethersulfone filter and a 5 μm glass fibre pre-filter inside a 50 mm polypropylene capsule with luer lock fitting connections. Water was filtered and the filter air-dried using a Bürkle Vampire Sampler (a peristaltic pump with a flow rate of up to 2 L per minute) and a piece of silicone hose (1 m) with luer lock adapter. The hose was placed inside the sampling bag, filled with sample water, then the filter capsule was attached to the adapter before continuing to run the pump. Once all water had been filtered or the filter clogged, the hose was removed from the sampling bag and pump run until no more water exited the filter. The filter capsule was detached from the hose, following which the capsule was filled with 1.5 mL of Longmire’s solution for DNA preservation (Longmire et al., 1997). A new kit and a new piece of silicone hose (1 m) with luer lock adapter were used for each pond. Field blanks (one of each kit type using mineral water instead of pond water) were included from May onwards to assess potential field contamination risk.

**Figure 2.**
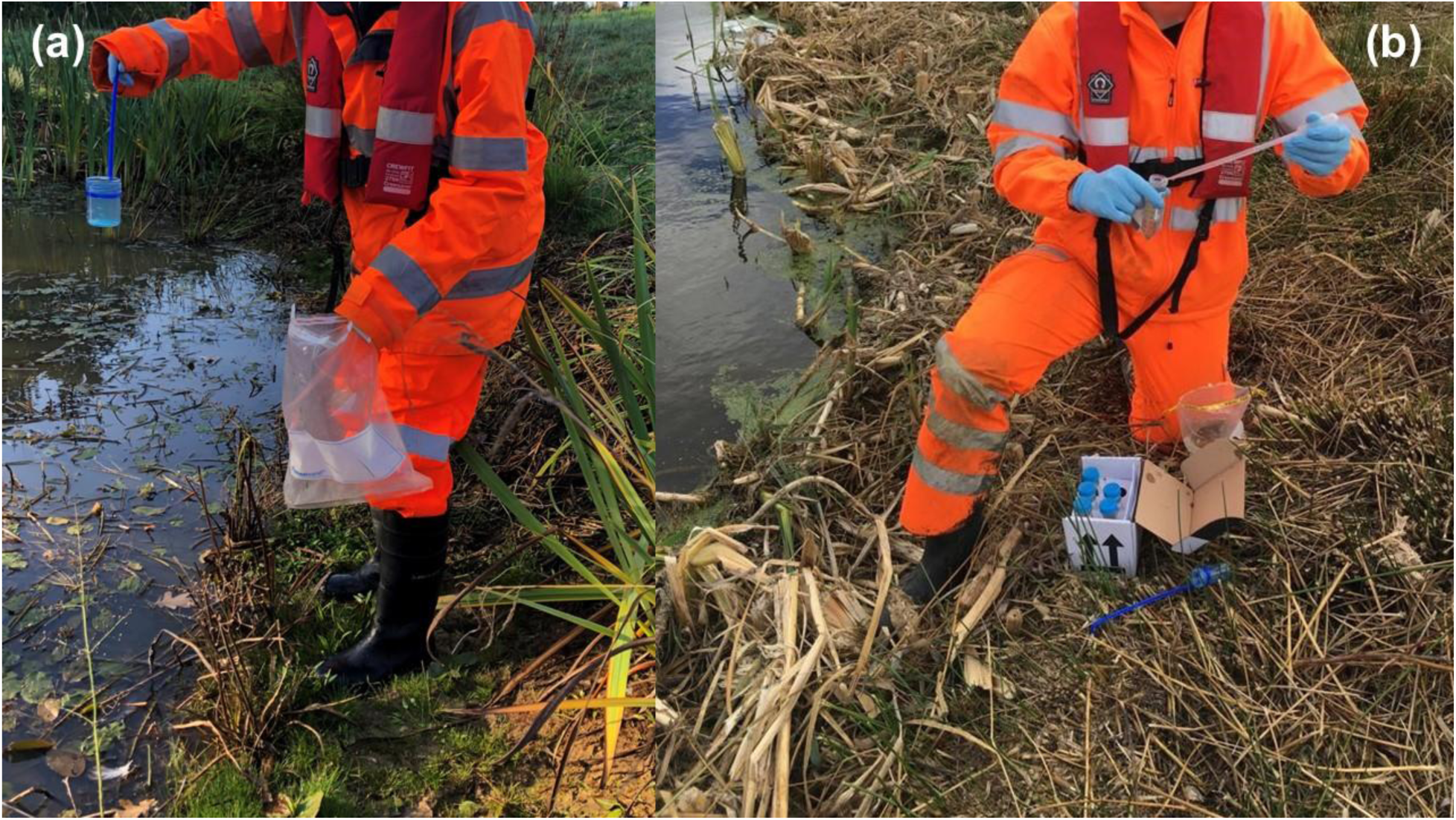
AtkinsRéalis ecologists **(a)** collecting water for filtration and **(b)** processing water using an EP kit.

The water sampling protocol established by Biggs et al. (2014) was used for both eDNA capture methods with minor modifications for filtration. For EP, 20 x 30 mL subsamples were collected at equidistant intervals around the pond perimeter then pooled into a single 900 mL sampling bag for homogenisation, following which 6 x 15 mL aliquots were added to tubes containing 33 mL of absolute ethanol and 1.5 mL of sodium acetate (3 M). For filtration, 20 x 125 mL subsamples were collected at equidistant intervals around the pond perimeter and pooled into a single 3.5 L sampling bag for homogenisation, following which as much water as possible was filtered. For EP, 600 mL of water was always collected, and 90 mL invariably processed. For filtration, 2.5 L of water was always collected, with up to 2.5 L filtered but volumes varied for each pond in each month (Supporting Information: Appendix 2, Figure S3). Water sampling for EP or filtration took the same amount of time (20-30 minutes), but water processing was typically faster with the EP kit (∼10 minutes) than the filtration kit (∼20 minutes) due to the greater volumes of water being processed with filtration (Supporting Information: Appendix 2, Figure S3). EP samples were stored at ambient temperature for same-day courier collection or refrigerated until courier collection. Filtration samples were stored at ambient temperature.

Where possible, population size class assessment surveys were performed for each pond that was positive for great crested newt prior to 2022 (i.e. generally based on surveys within the last four years for most ponds but occasionally based on surveys dating back to 2013 for some ponds). These were based on six visits from May to June and one visit in October using different combinations of bottle trapping, torchlight survey, refuge searching and netting. Bottle trapping could not be used where protected mammal species were present and torchlight could not be used for ponds with extensive vegetation cover (Table 1). Further details on each of these survey methods can be found in Langton et al. (2001). Pond 25 was subject to an out-of-season population survey only as this pond was included from July onwards as a replacement for an eDNA positive pond which became dry. Ponds 8, 9, 17, 21 and 22 were subject to an in-season population survey only as these ponds had dried out and/or become unsafe to access by October. Ponds 1-4 and 24 were not assessed as these were control ponds known or suspected to be negative for great crested newt, and Ponds 18, 19 and 23 were not assessed as they dried before population survey could occur.

### 2.3. Laboratory analysis

At NatureMetrics, resulting pellets from EP kits and filters from filtration kits were extracted using Qiagen DNeasy Blood and Tissue kits following the manufacturer protocol, with elution in 200 µL of Buffer AE. For EP kits, eDNA was precipitated via centrifugation at 14,000 x g and processed in accordance with Biggs et al. (2014). Briefly, after centrifugation, supernatant from all six subsamples was discarded, following which 360 µL of Buffer ATL was added to one tube for vortexing before the supernatant was transferred to the next tube. The supernatant in the sixth tube, containing DNA from all six subsamples, was transferred to a 2 mL tube for addition of Proteinase K (20 µL). DNA extraction then followed manufacturer protocol. For filtration kits, eDNA was extracted following a modified version of the Spens et al. (2017) SX capsule method for disc filters in buffer, with Proteinase K added directly to the filter housing to minimise the risk of contamination arising from handling of the filter. After incubation, the lysate was removed from the filter housing using a luer lock syringe and added to a 2 mL tube for DNA extraction following manufacturer protocol.

All samples were tested for degradation and inhibition using two commercial quantitative PCR (qPCR) kits that target two separate synthetic Internal Positive Controls (IPC; Eurogentec). One IPC was included in the ethanol and sodium acetate of EP kits or the preservative solution of filtration kits to test for degradation, and one IPC was added to DNA extracts to test for inhibition. qPCR amplification for each IPC was carried out in duplicate on each sample. If at least one of the replicates showed a shift of greater than two Ct values relative to controls, the sample was considered degraded and/or inhibited (depending on IPC). Inhibited samples were diluted 1:2 and 1:4 before species-specific qPCR amplification in accordance with Biggs et al. (2014). qPCR amplification was then carried out in 12 technical replicates per sample using species-specific primers and probe for the great crested newt, developed by Thomsen et al. (2012) and adopted by Biggs et al. (2014), in the presence of extraction negative controls, qPCR positive controls, and qPCR negative controls. The great crested newt qPCR reagents (and their concentrations and volumes) and conditions were as specified in the WC1067 protocol (Biggs et al., 2014), with 3 µL of template DNA used in each reaction. A score is given for the number of positive technical replicates out of 12, i.e. eDNA score. A sample is deemed positive for great crested newts if the eDNA score is 1 or higher. Results from the great crested newt qPCR assay are considered to have a high confidence rating according to the Confidence Assessment Tool for eDNA qPCR Results (K. J. Harper et al., 2021). Quality control methods included the use of field blanks, kit blanks, extraction negative and qPCR negative controls, and qPCR positive controls. The qPCR positive controls are standards of known great crested newt DNA concentration amplified in triplicate to determine correct amplifications.

### 2.4. Data analysis

Data were analysed in R v4.2.2 (R Core Team, 2022), and all plots were produced using the packages mapview v2.11.2 (Appelhans et al., 2023), leafsync v0.1.0 (Appelhans & Russell, 2019), ggplot2 v3.5.1 (Wickham, 2016), ggpubr v0.6.0 (Kassambara, 2020), and patchwork v1.3.0 (Pedersen, 2024). Control ponds (ponds 1-4 and 24) and negative process controls were assessed for any evidence of contamination, then these were removed from the data set.

#### 2.4.1. Generalised Linear Mixed-effect Models

Great crested newt detection success (both detection rate and eDNA scores) with EP and filtration in each month was statistically compared for Ponds 5-23 and 25. Next, for ponds with both in-season (April to June) and out-of-season (July to October) data (Ponds 5-16, 20 and 22), months were grouped as in-season or out-of-season to statistically compare detection success with EP and filtration between these two time periods. We examined the influence that September and October results had on in-season vs. out-of-season detection by repeating the statistical comparison without these data, i.e. only July and August results grouped as out-of-season (see Supporting Information for results of in-season vs. out-of-season comparisons).

Generalised Linear Mixed-effects Models (GLMMs) with binomial error family were used for these analyses. eDNA scores were converted to proportional eDNA scores (i.e. ranging from 0 to 1) for modelling. These GLMMs included an interaction term between eDNA capture method and month or time period, but the interaction term was dropped if it was not significant. eDNA capture method and month or time period were retained as fixed effects. We also examined the influence of volume of water filtered on proportional eDNA scores using a GLMM with binomial error family (see Supporting Information).

In-season population sizes for Ponds 5-17 and 20-22 were available as quantitative (peak adult counts) or qualitative (class) measurements. Classes were ‘small’ (up to 10 animals counted), ‘medium’ (11–100 animals counted) and ‘high’ (counts exceeding 100 animals) (English Nature, 2001). The population size and eDNA data for these ponds were split by eDNA capture method. The EP and filtration subdatasets were each analysed using three GLMMs with binomial error family, where both population size measurements were included as fixed effects. The three GLMMs had different response variables: (1) corresponding eDNA score, i.e. proportional eDNA score corresponding to May when most in-season survey visits for population size assessment were made; (2) average eDNA score, i.e. the average of proportional eDNA scores for in-season months; and (3) peak eDNA score, i.e. the highest proportional eDNA score observed across in-season months.

Pond number was included as a random effect in all GLMMs to account for repeat visitation and unknown variation inherent to individual ponds. GLMMs, model performance checks, extraction of model predictions, and post hoc analyses were implemented using the packages glmmTMB v1.1.10 (Brooks et al., 2017), DHARMa v0.4.7 (Hartig, 2022), easystats v0.7.3 (Lüdecke et al., 2022), arm v1.14-4 (Gelman & Su, 2022), emmeans v1.10.5 (Lenth, 2022), and effects v4.2-2 (Fox & Weisberg, 2018; Fox & Weisberg, 2019). If a significant effect was found for variables using the *drop1* function from the package stats v4.2.2 (R Core Team, 2022), multiple pairwise comparisons were performed using the *pairs* function from the package emmeans v1.10.5. The full data set has been provided as Supporting Information: Appendix 2.

#### 2.4.2. Occupancy models

A potential flaw with GLMMs is they do not account for imperfect detection via a hierarchical dependence structure like multi-scale occupancy models do. Like other monitoring methods, eDNA surveys do not always detect a species when it is present, i.e. false negative. The difference with eDNA surveys is false negatives can occur at different stages: species may not release enough DNA into the environment, eDNA may not be captured during sample collection, and/or eDNA may not be amplified during laboratory analysis (Dorazio & Erickson, 2018). Standard occupancy models assume no false positives, i.e. species is absent but recorded as present. This can occur with other survey methods where a surveyor mistakenly identifies a species based on visual or audio cues, but may be more pronounced with eDNA surveys where a species’ eDNA can be transported to sites where the species is not present by surveyors or other wildlife, or contamination can occur in the laboratory (Griffin et al., 2020).

There are now several multi-scale occupancy models available for eDNA data, which account for false negative (and occasionally false positive) error in the field and the laboratory. However, each has advantages and disadvantages, including the type of error that can be estimated, the ability to specify different covariates at different levels, and the ability to account for repeat visits to the same sites using multiple survey methods. We applied three of these models to our data to examine false negative and false positive error rates across months and with different eDNA capture methods.

The Griffin et al. (2020) model, implemented via the package eDNAShinyApp v1.0 (Diana et al., 2021), enables false positive and false negative error rates to be estimated for eDNA surveys. This allows specification of covariates which may influence site occupancy and detection probability, but different covariates cannot be specified at different levels. In our case, we would expect eDNA capture method and month to influence eDNA detection probability in water samples (Stage 1, θ), but we would only expect eDNA capture method to influence eDNA detection probability in qPCR replicates (Stage 2, *p*) due to differences in DNA extraction. The Dorazio and Erickson (2018) model, implemented with a Gibbs sampler via the package msocc v1.1.0 for computational efficiency (Stratton et al., 2020), does allow different covariates to be specified at different levels, but only provides estimates of false negative error. Neither of these models account for repeat visits to the same site in the same way as a GLMM with a random effect does. Instead, each visit is treated as a unique site. Multi-method models (Nichols et al., 2008), implemented via the package RPresence v2.15.19 (MacKenzie & Hines, 2025), account for non-independence between sampling methods at shared, local scale sampling locations (Nichols et al., 2008). However, these implement a different hierarchical dependence structure (where Ψ is occupancy at landscape scale, θ is local-scale occupancy, and *p* is probability of detection) and only estimate false negative error rate.

Specification of each model is described in the Supporting Information. We present the results of these models but note that none are particularly well-suited to our experimental design, i.e. multiple eDNA sampling methods used on repeat visits to the same sites.

## 3. Results

### 3.1. Literature review

Of seven studies identified, only two published studies used filtration for eDNA capture, where up to 1 L of water was processed. Across these two studies, filtration recovered more great crested newt eDNA than EP in a tank experiment but not in natural ponds where eDNA concentration was similar between methods (Buxton et al., 2017; 2018b). The other five studies (two published and three grey literature) used EP and demonstrated that out-of-season detection was possible, including a 2019 data set supplied by AtkinsRéalis where every pond surveyed was positive for great crested newt eDNA in September (Gorman et al., 2020). There were differences in sampling methodology across all studies, but the eDNA capture conditions for EP and filtration, the DNA extraction method, and the amplification method were consistent (see Supporting Information: Appendix 1).

### 3.2. Field study

#### 3.2.1. Quality control

Results of quality control checks are summarised in Figure 3 and Supporting Information: Appendix 2. Of the five control ponds (Ponds 1-4 and 24), three were still negative in 2022 (ponds 1-3) but the other two (ponds 4 and 24) produced positive results from filters. Pond 4 (eDNA score = 1) was positive in June, and Pond 24 was positive in October (eDNA score = 2). All samples passed tests for degradation and inhibition except four filters in May (Pond 22), September (Ponds 3 and 4) and October (Pond 24) which were inhibited despite troubleshooting undertaken to resolve inhibition. Samples are analysed for great crested newt regardless of inhibition and degradation being identified. Results from the two filters for Ponds 22 and 24 were returned as positive rather than inconclusive because great crested newt DNA amplified successfully, and therefore these samples were retained. However, results for the two filters from Ponds 3 and 4 were returned as inconclusive rather than negative because great crested newt DNA did not amplify which could be due to inhibition or absence of great crested newt DNA in the sample. One field blank (EP kit) also produced an inconclusive result in October because it failed the inhibition test. All other negative process controls were negative for great crested newt DNA and passed quality control.

**Figure 3.**
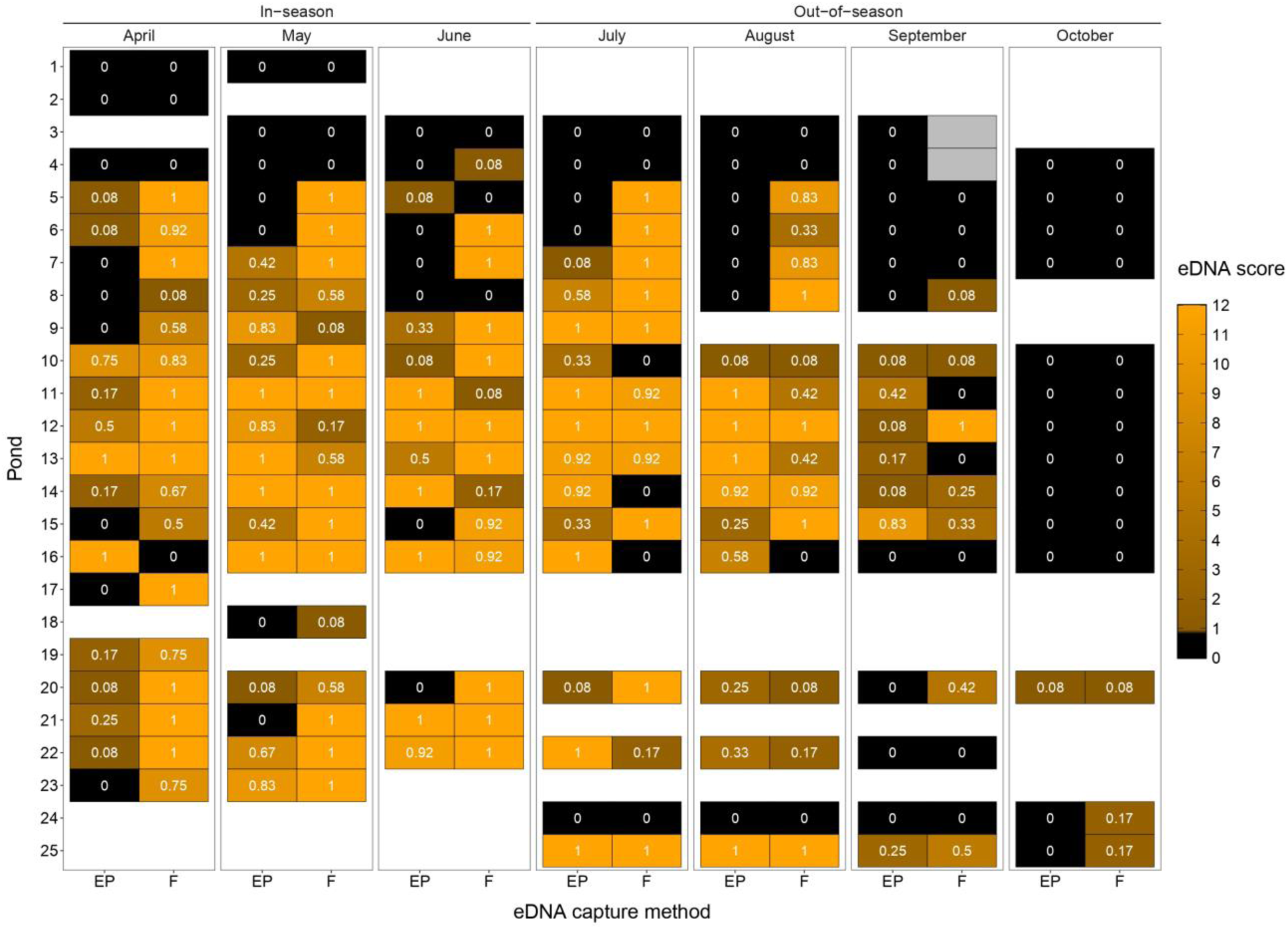
Great crested newt status of ponds surveyed from April to October using ethanol precipitation (EP) kits and filtration (F) kits. A pond is classed as positive for positive for great crested newts if one or more of the qPCR replicates performed on the eDNA sample are positive, i.e. eDNA score >= 1. Cells are shaded according to kit result (black = negative, orange = positive, grey = inconclusive), with positive results on a colour gradient according to eDNA score. White cells represent months where a pond was not surveyed due to drying out and/or health and safety concerns. Numbers in cells indicate proportional eDNA scores used for modelling.

#### 3.2.2. Sample size

In total, 25 ponds were sampled from April to October (Table 1, Figure 3). Ponds 1 and 2 were newly created drainage ponds and produced negative eDNA results in 2022. Pond 3 was not previously found to support great crested newts and also produced negative eDNA results in 2022. Ponds 4 and 24 were negative for great crested newts until 2022 when they produced positive eDNA results. The remaining 20 ponds that were previously found to support great crested newts also produced positive eDNA results in 2022 (Figure 3; Supporting Information: Appendix 2).

Of the 25 ponds sampled that produced positive and negative results over the course of the study, 16 had in-season and out-of-season eDNA data, while seven had in-season eDNA data only (typically spanning 2 months) and two had out-of-season eDNA data only. Two of the 16 ponds with both in-season and out-of-season eDNA data were control ponds. This left 14 ponds that were previously positive for great crested newts using conventional and eDNA surveys that also produced a positive eDNA result for at least one in-season month in 2022. All 14 ponds were surveyed in July, decreasing to 13 ponds in August and September, and 11 ponds in October due to ponds drying out and/or health and safety issues (Figure 3).

All control ponds and laboratory controls were excluded from downstream analysis. All remaining ponds (Ponds 5-23 and 25, n = 20) were all used in statistical comparisons of eDNA detection success in each month (sections 3.2.3 and 3.2.5). Great crested newt ponds with both in-season and out-of-season eDNA data (Ponds 5-16, 20 and 22, n = 14) were used in statistical comparisons of eDNA detection success in-season vs. out-of-season (Supporting Information). Great crested newt ponds with in-season eDNA and population data (Ponds 5-17 and 20-22, n = 16) were used in statistical analysis of eDNA score in relation to population size (section 3.2.4).

#### 3.2.3. eDNA capture method and month

The number of positive ponds generated by each eDNA capture method in each month ranged from 1 to 17 (Figure 4a). The effect of eDNA capture method did not depend on month and vice versa as the interaction term was not significant (Χ^2^_6_ = 9.101, P = 0.168), but the individual effects of eDNA capture method (Χ^2^_1_ = 8.030, P = 0.005) and month (Χ^2^_6_ = 61.959, P < 0.001) were significant (Table 2). Filtration (mean probability of a positive eDNA result ± SE = 0.823 ± 0.057) consistently produced more positive ponds than EP (0.608 ± 0.083) overall (z = -2.727, P = 0.006). May (0.924 ± 0.045) had the highest number of positive ponds and significantly differed to September (0.484 ± 0.124, z = 3.382, P = 0.013) and October (0.075 ± 0.052, z = 5.039, P < 0.001) but not other months (April and June to August: z = -0.969-1.353, P > 0.05). October had the fewest positive ponds and significantly differed to all other months (April to August: z = 4.628-5.039, P < 0.001), although the difference between September and October was on the cusp of significance (z = 2.981, P = 0.045; Figure 4b).

**Figure 4.**
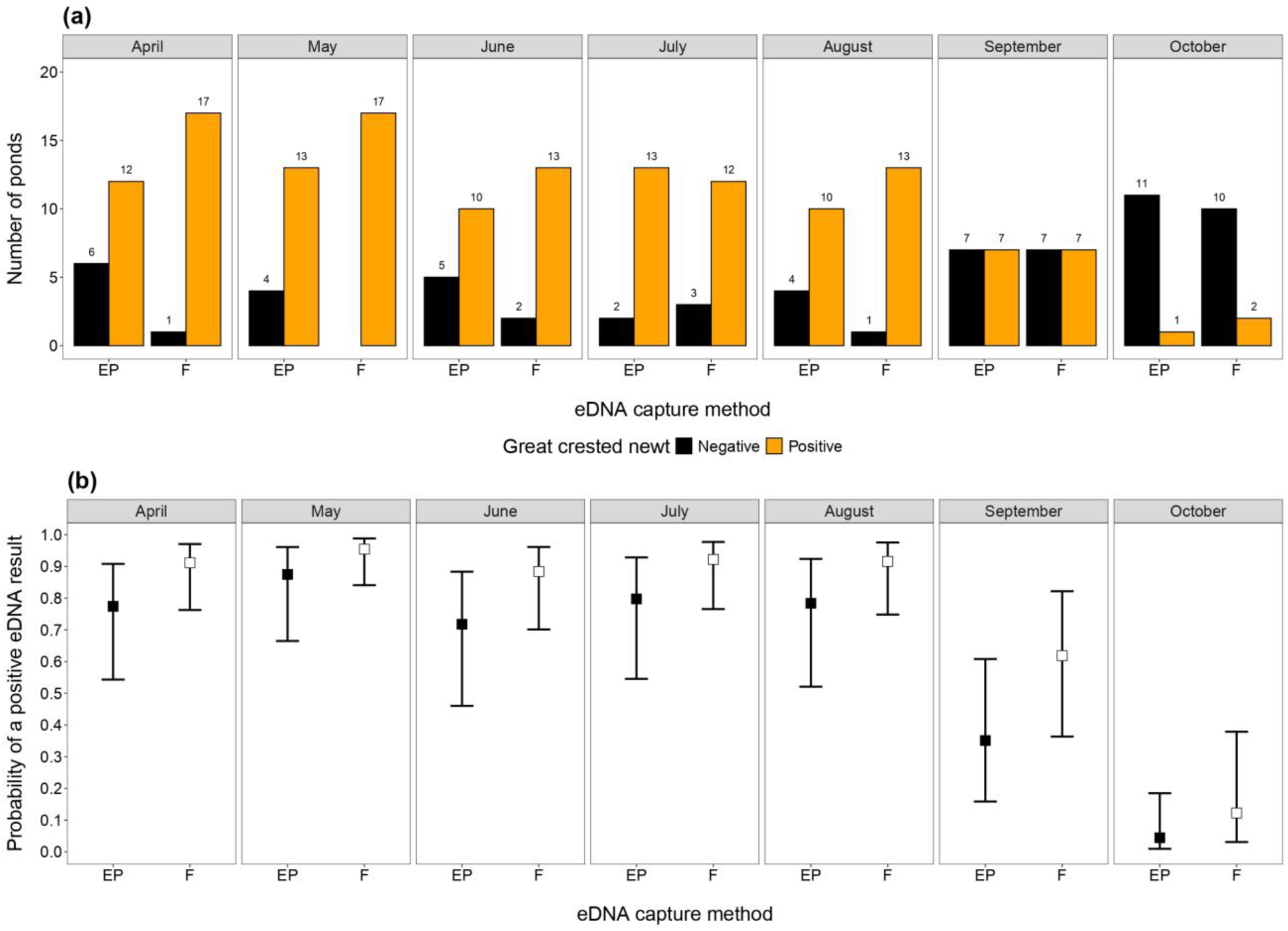
The effects of eDNA capture method and month on GCN detection. In **(a)**, bars show the number of ponds that were positive (orange) and negative (black) for great crested newt each month with EP and filtration (F) kits. The number of ponds corresponding to each category are displayed above bars. In **(b)**, square points represent the model-predicted positive eDNA result probabilities with EP (black) and F (white) in each month, and error bars show the 95% confidence intervals for these predictions.

**Table 2.** Summary of GLMM and post hoc analysis performed to examine the influence of eDNA capture method and month (as individual and interacting effects) on the GCN status of ponds. Significant p-values (<0.05) are in bold and df denotes degrees of freedom.

| GLMM |  |  |  |  | Post hoc analysis |  |  |  |  |
| --- | --- | --- | --- | --- | --- | --- | --- | --- | --- |
| Formula | Variable | Effect | | | Levels | Mean probability ± SE | Pairwise comparison | | |
|  |  | X<sup>2</sup> | df | P |  |  | Pair | z | P |
| GCN status ~<br>eDNA capture<br>method + month<br>+ eDNA capture<br>method : month | eDNA capture method : month | 9.101 | 6 | 0.168 | NA | NA | NA | NA | NA |
| | eDNA capture method | 8.030 | 1 | **0.005** | EP | 0.608 ± 0.083 | EP : F | -2.727 | **0.006** |
| | | | | | F | 0.823 ± 0.057 | | | |
| | month | 61.959 | 6 | **<0.001** | Apr | 0.856 ± 0.066 | Apr : May | -0.969 | 0.961 |
|  |  |  |  |  |  |  | Apr : Jun | 0.449 | 0.999 |
|  |  |  |  |  |  |  | Apr : Jul | -0.197 | 1.000 |
|  |  |  |  |  |  |  | Apr : Aug | -0.079 | 1.000 |
|  |  |  |  |  |  |  | Apr : Sep | 2.783 | 0.079 |
|  |  |  |  |  |  |  | Apr : Oct | 4.710 | **<0.001** |
|  |  |  |  |  | May | 0.924 ± 0.045 | May : Jun | 1.353 | 0.826 |
|  |  |  |  |  |  |  | May : Jul | 0.729 | 0.991 |
|  |  |  |  |  |  |  | May : Aug | 0.829 | 0.982 |
|  |  |  |  |  |  |  | May : Sep | 3.382 | **0.013** |
|  |  |  |  |  |  |  | May : Oct | 5.039 | **<0.001** |
|  |  |  |  |  | Jun | 0.815 ± 0.082 | Jun : Jul | -0.615 | 0.996 |
|  |  |  |  |  |  |  | Jun : Aug | -0.493 | 0.999 |
|  |  |  |  |  |  |  | Jun : Sep | 2.343 | 0.224 |
|  |  |  |  |  |  |  | Jun : Oct | 4.427 | **<0.001** |
|  |  |  |  |  | Jul | 0.872 ± 0.067 | Jul : Aug | 0.110 | 1.000 |
|  |  |  |  |  |  |  | Jul : Sep | 2.821 | 0.071 |
|  |  |  |  |  |  |  | Jul : Oct | 4.727 | **<0.001** |
|  |  |  |  |  | Aug | 0.863 ± 0.072 | Aug : Sep | 2.691 | 0.101 |
|  |  |  |  |  |  |  | Aug : Oct | 4.628 | **<0.001** |
|  |  |  |  |  | Sep | 0.484 ± 0.124 | Sep : Oct | 2.981 | **0.045** |
|  |  |  |  |  | Oct | 0.075 ± 0.052 |  |  |  |

The eDNA score produced by each eDNA capture method in each month ranged from 0 to 12 (proportional eDNA score = 0-1; Figure 5a). The effect of eDNA capture method did not depend on month and vice versa as the interaction term was not significant (Χ^2^_6_ = 4.888, P = 0.558), but the individual effects of eDNA capture method (Χ^2^_1_ = 12.903, P < 0.001) and month (Χ^2^_6_ = 52.380, P < 0.001) were significant (Table 3). Filtration (mean proportional eDNA score ± SE = 0.492 ± 0.085) produced higher eDNA scores than EP (0.233 ± 0.063) overall (z = -3.474, P < 0.001; Figure 5b). The highest eDNA scores were observed in July (0.691 ± 0.090) and lowest eDNA scores observed in September (0.143 ± 0.067) and October (0.011 ± 0.019). Scores from April to August did not significantly differ (April to August: z = -1.384-1.277, P > 0.05). Scores in May (0.654 ± 0.088), June (0.613 ± 0.096) and July significantly differed to scores in September (May: z = 3.638, P = 0.005; June: z = 3.343, P = 0.015; July: z = 3.801, P = 0.003), but not October (Table 3; Figure 5c).

**Figure 5.**
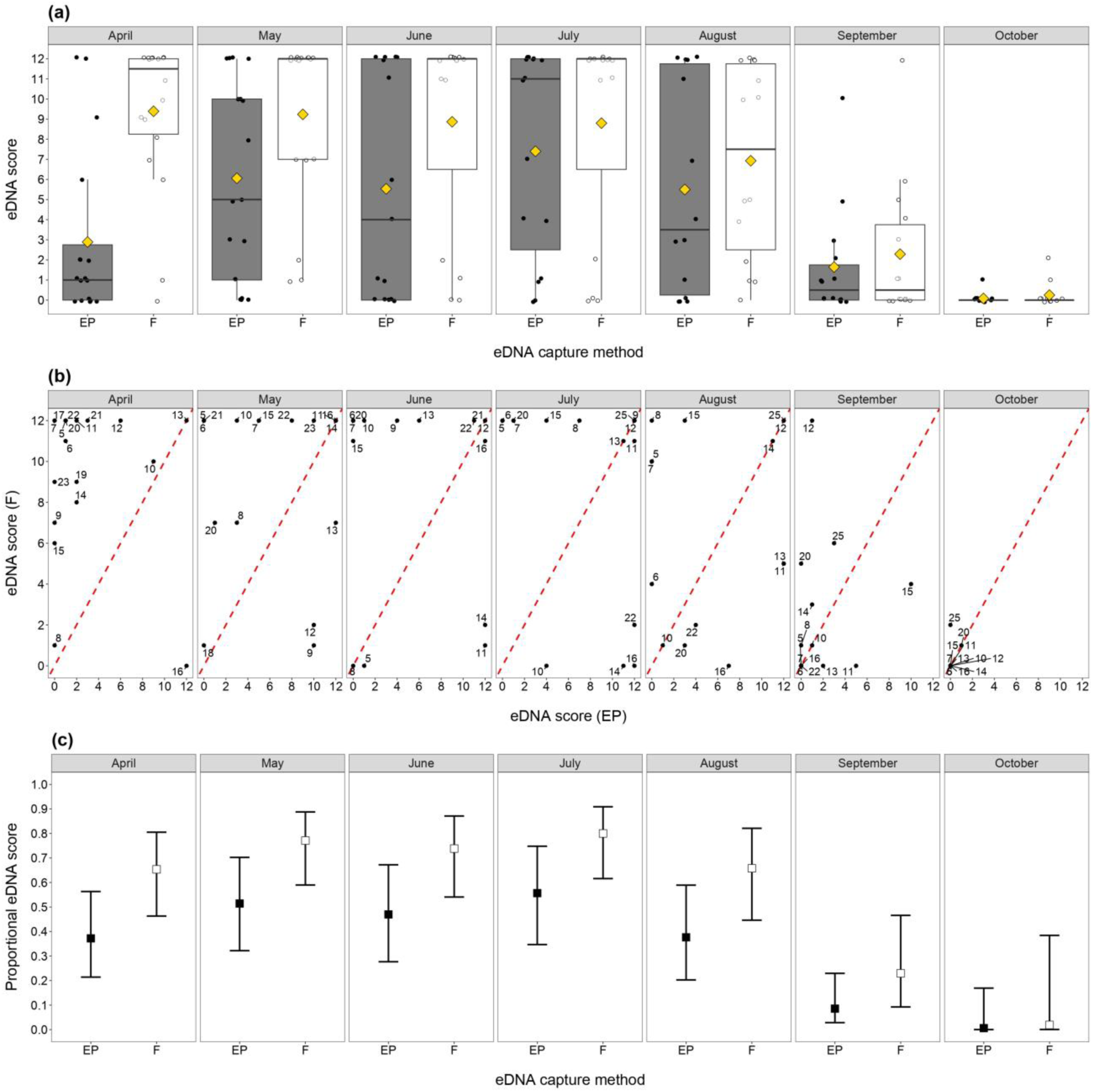
The effects of eDNA capture method and month on eDNA score. In **(a)**, the spread of eDNA scores with EP and filtration (F) kits in each month is represented by boxes which show the 25th, 50th and 75th percentiles, whiskers which show the 5th and 95th percentiles, and thick horizontal lines which show the median. Circular points (EP = black, F = white) represent the eDNA score for individual ponds, and yellow diamonds represent the mean eDNA score for EP and F. In **(b)**, eDNA scores with EP are plotted against eDNA scores with F for each pond in each month. The red dashed line represents the same eDNA score being produced by both eDNA capture methods; points to the left of this line are ponds where F produced a higher eDNA eDNA score and points to the right of this line are ponds where EP produced a higher eDNA score. Numbers represent the number allocated to each pond for the present study (see Table 1). In **(c)**, the square points represent the model-predicted proportional eDNA scores with EP (black) and F (white) in each month, and error bars show the 95% confidence intervals for these predictions

**Table 3.** Summary of GLMM and post hoc analysis performed to examine the influence of eDNA capture methods and month (as individual and interacting effects) on the proportional eDNA score of ponds. Significant p-values (<0.05) are in bold and df denotes degrees of freedom.

| GLMM |  |  |  |  | Post hoc analysis |  |  |  |  |
| --- | --- | --- | --- | --- | --- | --- | --- | --- | --- |
| Formula | Variable | Effect |  |  | Levels | Mean<br>proportional<br>eDNA score ±<br>SE | Pairwise comparison |  |  |
|  |  | X<sup>2</sup> | df | P |  |  | Pair | z | P |
| Proportional<br>eDNA score ~<br>eDNA capture<br>method +<br>month + eDNA<br>capture<br>method :<br>month | eDNA capture method : month | 4.888 | 6 | 0.558 | NA | NA | NA | NA | NA |
|  | eDNA capture method | 12.903 | 1 | <0.001 | EP | 0.233 ± 0.063 | EP : F | -3.474 | <0.001 |
|  |  |  |  |  | F | 0.492 ± 0.085 |  |  |  |
|  | month | 52.380 | 6 | <0.001 | Apr | 0.514 ± 0.090 | Apr : May | -1.122 | 0.922 |
|  |  |  |  |  |  |  | Apr : Jun | -0.760 | 0.989 |
|  |  |  |  |  |  |  | Apr : Jul | -1.384 | 0.811 |
|  |  |  |  |  |  |  | Apr : Aug | -0.034 | 1.000 |
|  |  |  |  |  |  |  | Apr : Sep | 2.861 | 0.064 |
|  |  |  |  |  |  |  | Apr : Oct | 2.535 | 0.147 |
|  |  |  |  |  | May | 0.654 ± 0.088 | May : Jun | 0.328 | 1.000 |
|  |  |  |  |  |  |  | May : Jul | -0.307 | 1.000 |
|  |  |  |  |  |  |  | May : Aug | 1.017 | 0.950 |
|  |  |  |  |  |  |  | May : Sep | 3.638 | **0.005** |
|  |  |  |  |  |  |  | May : Oct | 2.841 | 0.068 |
|  |  |  |  |  | Jun | 0.613 ± 0.096 | Jun : Jul | -0.615 | 0.996 |
|  |  |  |  |  |  |  | Jun : Aug | 0.682 | 0.994 |
|  |  |  |  |  |  |  | Jun : Sep | 3.343 | **0.015** |
|  |  |  |  |  |  |  | Jun : Oct | 2.741 | 0.088 |
|  |  |  |  |  | Jul | 0.691 ± 0.090 | Jul : Aug | 1.277 | 0.863 |
|  |  |  |  |  |  |  | Jul : Sep | 3.801 | **0.003** |
|  |  |  |  |  |  |  | Jul : Oct | 2.926 | 0.053 |
|  |  |  |  |  | Aug | 0.519 ± 0.102 | Aug : Sep | 2.788 | 0.078 |
|  |  |  |  |  |  |  | Aug : Oct | 2.533 | 0.147 |
|  |  |  |  |  | Sep | 0.143 ± 0.067 | Sep : Oct | 1.483 | 0.755 |
|  |  |  |  |  | Oct | 0.011 ± 0.019 |  |  |  |

#### 3.2.4. Population size

No great crested newts were found during population size class assessment surveys conducted in October, but three ponds produced a positive eDNA result. Pond 20 was positive using both filtration and EP kits, whereas Ponds 24 and 25 were only positive using filtration kits. Consequently, no analysis of population size in relation to eDNA score out-of-season was undertaken.

Sixteen ponds (Ponds 5-17 and 20-22) had in-season population size data (peak adult counts and classes) that could be used to analyse population size in relation to eDNA score. The remaining nine ponds were unsuitable because: a) they were control ponds, b) population size class assessment surveys were not undertaken as ponds were removed from the study due to drying out and/or health and safety reasons, or c) they were replacements added to the study from July onwards and only had out-of-season data.

There was no influence of peak adult counts or population class on corresponding eDNA score, average eDNA score, or peak eDNA score with EP or filtration (see Supporting Information: Table S3, Figure S4). Therefore, adult population size did not appear to influence the amount of great crested newt eDNA captured with either eDNA capture method.

#### 3.2.5. Occupancy modelling

With the Griffin et al. (2020) model implemented in the eDNAShinyApp package, the posterior mean occupancy for all sites was 0.078 with posterior credible intervals (PCI) ranging from 0.002 to 0.763. At Stage 1 (sampling), the true positive rate (mean posterior probability of θ_11_) was high, ranging from 0.556-0.711 (mean = 0.637; median = 0.628; PCI = 0.002-0.999954). The false positive rate (mean posterior probability of θ_10_) was moderate, ranging from 0.022-0.781 (mean = 0.426; median = 0.465; PCI = 0.001-0.949) (Figure 6ai). At Stage 2 (laboratory analysis), the true positive rate (mean posterior probability of *p*_11_) was very high at 0.936 (PCI = 0.918-0.952). The false positive rate (mean posterior probability of *p*_11_) was very low at 0.097 (PCI = 0.082- 0.118). False negative probability is calculated as 1—true positive: this was lower than the false positive rate at both Stage 1 (1 – 0.637 = 0.363), and Stage 2 (1 – 0.936 = 0.064). Both eDNA capture method and month were found to have Posterior Inclusion Probability (PIP) scores of 1 for Stage 1 true positive and false positive rates, suggesting an influence on probability of great crested newt eDNA capture in water samples. The probability of true positives at Stage 1 was highest in July, followed by May then June. Filtration also produced a higher probability of true positives. However, these patterns were the same for false positives (Figure 6aii).

With the Dorazio & Erickson (2018) model implemented in the msocc package, the model with eDNA capture method and month as sample-level covariates (but no interaction between these), and eDNA capture method as a replicate-level covariate performed best. The probability of eDNA capture in a water sample (θ) was higher with filtration each month (median values: April = 0.874, May = 0.934, June = 0.856, July = 0.902, August = 0.895, September = 0.627, October = 0.182) than with EP (median values: April = 0.723, May = 0.845, June = 0.695, July = 0.778, August = 0.762, September = 0.390, October = 0.078), albeit with overlapping CI. The probability of eDNA capture in a water sample was highest in May then July and August with either EP or filtration, and lowest in October (Figure 6bi). Detection probability in a qPCR replicate (*p*) was higher with filtration (0.744) than EP (0.568), with no overlapping CI (Figure 6bii).

With the Nichols et al. (2008) model implemented in the RPresence package, the model included eDNA capture method and month as covariates for detection (*p*). Filtration had a higher predicted probability of detection than EP in most months except July. Probability of detection was highest in May, followed by April then August with filtration, or July followed by May then August with EP. Probability of detection was lowest in October with either eDNA capture method (Figure 6c).

**Figure 6.**
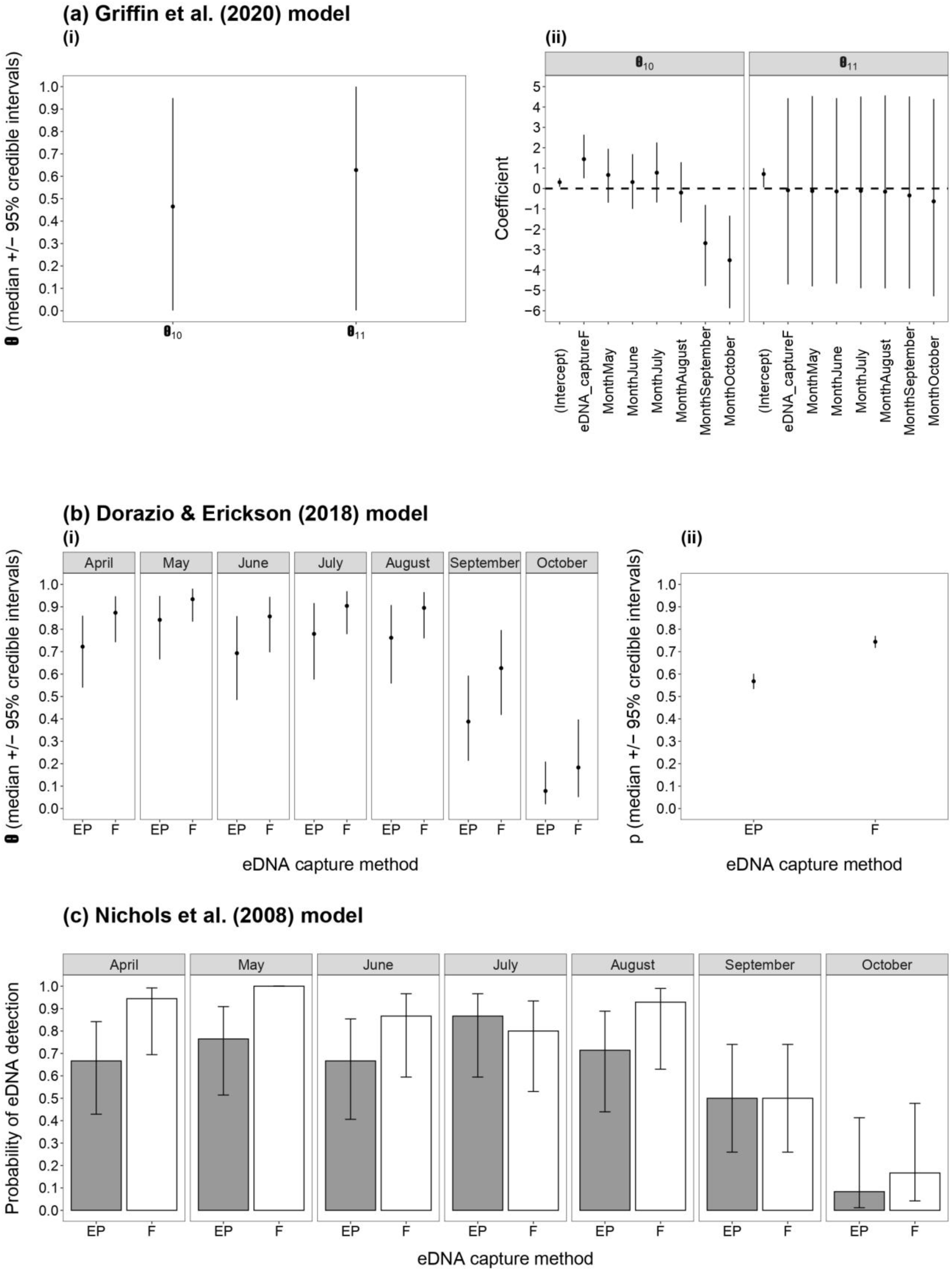
Outputs of three different multi-scale occupancy models for levels at which covariates were considered. For the Griffin et al. (2020) model **(a)**, the θ_11_ (true positive error rate) and θ_10_ (false positive error rate) estimates are shown with posterior credible intervals **(i)** alongside the Stage 1 coefficients and posterior credible intervals for each level of covariates where PIP values were greater than 0.5 **(ii)**. For the Dorazio & Erickson (2018) model **(b)**, the MCMC estimates of the parameters θ as a function of eDNA capture method (EP or filtration – F) and month **(i)**, and *p* as a function of eDNA capture method **(ii)** are shown, summarised as the median and 95% credible intervals of estimated values. For the Nichols et al. (2018) model **(c)**, the parameter estimates for *p* with EP and filtration (F) in each month are shown with 95% credible intervals.

The results of all three multi-scale occupancy models are consistent with the GLMM analysis, where filtration improved the probability of eDNA capture and subsequently great crested newt detection, with similar detection probabilities from April to August. However, the Griffin et al. (2020) model does indicate some potential for false positives.

## 4. Discussion

### 4.1. Quality control

Filtration for two control ponds produced unexpected positive eDNA results. Pond 24 is adjacent to a pond (∼50 m apart) not sampled in this study where great crested newt presence was confirmed in 2020, thus this positive detection may have resulted from great crested newts migrating from the adjacent pond in combination with higher sensitivity of filtration due to larger volumes of water processed (Rees et al., 2023). Pond 4 is located on a site where all ponds were likely or suspected to be negative for great crested newts and produced negative results either side of the positive result in June. Field and laboratory contamination is unlikely as all negative process controls were negative for great crested newt DNA. Possible explanations for sporadic eDNA detection include great crested newt adults or sub-adults temporarily using ponds for foraging or aquatic refuge (Priol et al., 2023), or eDNA transport by humans, domestic animals or other wildlife, e.g. great crested newt eDNA in water trapped in waterfowl feathers or predated great crested newts in waterfowl faeces (Beng & Corlett, 2020). At each of the three sites, it is not possible to rule out great crested newt dispersal from unsampled ponds in the wider landscape. It is also not possible to eliminate hydrological movement at the Aylesbury site where ponds are in closer proximity to each other and a stream is present (Figure 1c).

One field blank (EP kit) failed the inhibition test. Reasons for this are unknown but could include chemicals or ions present in the mineral water or contamination with an inhibitor present in the field (Schrader et al., 2012). However, given that all other negative process controls were negative for great crested newt DNA, great crested newt DNA was unlikely to be present in this field blank either.

### 4.2. Great crested newt eDNA survey season

Both the desk and field studies indicated that great crested newts can be detected in spring as well as summer, likely due to the presence of adults and larvae (Buxton et al., 2017; Rees et al., 2017), with reduced detection from September onwards. Therefore, the survey window could potentially be doubled (c. 15 April – 31 August instead of 15 April – 30 June), which would facilitate land access and relieve logistical pressures for ecological consultants, commercial eDNA providers, and developers (Rees et al., 2017).

The great crested newt eDNA assay has undergone substantial validation, thus a positive eDNA result means that great crested newts are very likely present and a negative eDNA result means great crested newts are likely absent assuming *appropriate timing* and replication in sampling (Thalinger et al., 2021). Ponds that were positive with either EP or filtration in July and August in the present study are likely to be true positives as most eDNA scores were high (Figure 3). Extending the great crested newt eDNA survey season could enable identification of ponds used for breeding and occupied by larvae as well as ponds only used for foraging or refuge that are critical to great crested newt survival but may be missed by the current eDNA survey window (Buxton et al., 2021b; Priol et al., 2023). Out-of-season detections could be used to begin planning conservation or mitigation activities (Gorman et al., 2020), including which waterbodies should be subject to in-season survey the following year (Rees et al., 2023). However, out-of-season non-detections require more scrutiny and cautious interpretation (Buxton et al., 2021b).

Although positive eDNA results outside the sampling timeframe originally used in assay validation are reliable (in so far as that they have the same potential for error as positive results in the timeframe used for validation), negative eDNA results are not as the survey timing is no longer optimal for DNA release from the target species (Thalinger et al., 2021; Troth et al., 2021). False negatives were observed for three ponds out-of-season. In these cases, great crested newts were detected with either EP or filtration in one month, followed by a non-detection the next month, then detection in the next month. This anomaly was also observed across in-season months in 10 ponds. This highlights the influence of eDNA capture method and population dynamics on great crested newt eDNA detection. Great crested newts exist as spatially structured populations with many interconnected sub-populations within the wider landscape. Individuals are capable of frequent short-distance (<400 m) dispersal and occasionally long-distance (>1 km) dispersal between ponds. Some ponds are ‘sources’ (i.e. core breeding ponds) whereas others are ‘sinks’ (i.e. occupied but not primarily used for breeding), resulting in a patchy population or metapopulation structure depending on the direction of dispersal. Sources and sinks can change over time due to environmental conditions (Griffiths et al., 2010; Unglaub, Cayuela et al., 2021).

As the highest number of positive ponds with either eDNA capture method was observed in May, the peak month of the core breeding season, regulators need to consider what acceptable levels of uncertainty and error are if eDNA surveys are undertaken in different months of the breeding season or outside the breeding season.

Waterbodies occupied by breeding great crested newt adults, eggs or larvae may produce false negatives if: they support fewer individuals or were not sampled when breeding activity peaked (both of which increase the rate of eDNA release and amount of eDNA present); they were not utilised by great crested newts in a given year; all eggs and larvae were predated; or the pond temporarily dried then refilled. In our study, two ponds that dried by July or August were occupied by great crested newts from April to June or July, and four ponds were occupied in April and/or May before drying. More ponds are likely to experience temporary drying before refilling in winter with declines in summer precipitation and increases in summer temperature and winter precipitation (Griffiths et al., 2010). Therefore, variable use of ponds is a key consideration for out-of-season survey.

Waterbodies occupied by non-breeding great crested newt adults in spring only may not be identified by eDNA surveys in July and August, resulting in false negatives (Buxton et al., 2017, 2021b). Similarly, waterbodies without larvae and increasingly fewer great crested newt adults as they disperse during out-of-season months, or waterbodies that are sporadically used by great crested newt adults and sub-adults for foraging and refuge in out-of-season months, may produce false negatives if great crested newts are not present at densities that would produce sufficient amounts of eDNA for detection (Buxton et al., 2021b; Priol et al., 2023). Cruickshank et al. (2021) found the number of non-breeding amphibian populations in Switzerland was substantially higher than breeding populations. Non-breeding populations also had lower detection probabilities than breeding populations due to lower abundance. Extinction risk of non-breeding populations may be lowered through habitat management thereby increasing breeding probability. Consequently, identification of non-breeding populations is key in promoting the favourable conservation status of a species.

Out-of-season great crested newt survey is unlikely to be appropriate for determining species absence and is an important caveat attached to regulatory acceptance of out-of-season eDNA survey. Interpretation of non-detections out-of-season as great crested newts being absent poses a potential risk to conservation or mitigation under UK and European legislation (Buxton et al., 2021a, b). To account for this risk, waterbodies that produce a negative result out-of-season should also be resurveyed the following spring to confirm whether great crested newts are indeed present or absent.

Nonetheless, false negatives can be encountered with any monitoring tool (Buxton et al., 2021a, 2022) as evidenced by non-detection with torching, bottle trapping or refuge searches at three ponds in October which produced positive eDNA results albeit with low eDNA scores. While these eDNA detections could be false positives, they could be true positives that highlight the detection sensitivity of eDNA analysis for detecting ponds with low numbers of great crested newts, possibly facilitated by processing more water (Rees et al., 2023). Other studies have also shown eDNA detection via single species or metabarcoding assays outperformed conventional survey methods for amphibians (Moss et al., 2022; Wikston et al., 2023), including great crested newts (Biggs et al., 2014, 2015; L. R. Harper et al., 2018; Priol et al., 2023; Rees et al., 2014). Ideally, multiple survey methods and repeated sampling events should be used to minimise false negatives, including eDNA and conventional surveys where time and resources allow (Moss et al., 2022; Rees et al., 2014; Wikston et al., 2023). False negatives could also be minimised by increasing biological or field sample replication (Buxton et al., 2021a, 2022; L. R. Harper et al., 2018).

False positive detections, as potentially indicated by a low eDNA score (Buxton et al., 2022), could be verified by reanalysing samples (Rees et al., 2014), increasing the eDNA score threshold for classing a sample as positive (Buxton et al., 2021a), or applying statistical models that account for false negative and false positive error rate (Buxton et al., 2021a, 2021b, 2022; Griffin et al., 2020). The Griffin et al. (2020) model indicated a moderate portion of detections in our study could be false positives originating from field sample collection, whether due to contamination from surveyors, other wildlife or hydrological connectivity, yet no field negative controls returned unexpected results (see section 4.1). The eDNAShinyApp package was not well-equipped to handle our experimental design (i.e. repeat visits to the same ponds), and we had a small number of sites, so it is possible that the false positive rate observed is inflated. The development of a multi-scale occupancy model with a hierarchical dependence structure that can handle repeat visits using multiple methods for eDNA data would enable more accurate estimation of false positive and false negative error rates. In the interim, applying the Griffin et al. (2020) model to similar data sets involving unique sites may improve understanding of false positive and false negative error rates for different eDNA capture methods and survey dates. However, all of the measures described to handle false positives would require further modification to the approved great crested newt eDNA protocol (Biggs et al., 2014).

Critically, surveys could not be conducted in February and March 2022 due to the timeframe for funding proposal submission, review, and acceptance. However, great crested newt adults are known to start returning to waterbodies in these months and even begin breeding in February and March (Buxton et al., 2017; Rees et al., 2017), especially in colder climates (L. R. Harper et al., 2019). Compared to in-season months, Rees et al. (2017) observed lower eDNA scores in February but similar eDNA scores in March. Additional work is needed to confirm whether survey in these months produces detection rates comparable with April to August or lower detection rates like those seen in September and October in the present study and previous studies (Buxton et al., 2018a, b; Rees et al., 2017). Ideally, this would be contracted by regulators in a large- scale, multi-year evaluation of great crested newt eDNA detection in-season versus out- of-season (February to October), encompassing waterbodies of known great crested newt status and variable eDNA scores throughout the UK which span the full spectrum of physicochemical variability seen in these systems (Rees et al., 2017, 2023). However, local-scale evaluations (either independent or as part of regulatory monitoring) will also generate vital evidence that will further improve understanding of out-of-season great crested newt eDNA detection and allow regulators to make informed decisions.

### 4.3. eDNA capture method

EP and filtration were comparable in terms of eDNA yield and eDNA score in the published studies that have compared both eDNA capture methods for great crested newt DNA to date (Buxton et al., 2017, 2018b; Rees et al., 2024). In the present field study, we found that filtration outperformed EP, both in terms of number of positive ponds and eDNA score, echoing other published work where filtration outperformed EP for targeted eDNA analysis (e.g. Hinlo et al., 2017; Minamoto et al., 2015; Peixoto et al., 2020; Spens et al., 2017; Troth et al., 2020).

Importantly, there were key differences between our study and previous works. Buxton et al. (2017, 2018b) surveyed eight small ponds (600 L, 1 m by 2 m and up to 0.6 m deep) within a small plot of land owned by the University of Kent. A single 1 L surface water sample was taken from the centre of each pond, thus the field protocol differed to Biggs et al. (2014). Glass microfibre (0.7 μm) filters were used and frozen until DNA extraction. More recently, Rees et al. (2024) analysed polyvinylidene difluoride (0.45 μm) filters from 88 natural ponds across Cheshire, Yorkshire and Norfolk, with filtered volumes ranging from 25 mL to a maximum of 500 mL. An ethanol-based preservative solution was added to filters for storage at ambient temperature for up to 8 weeks prior to DNA extraction. In our study, ponds ranged in surface area from 2 to 2,770 m^2^ (mean = 238.68 m^2^, median = 80 m^2^; Table 1), and the field protocol from Biggs et al. (2014) was adapted to accommodate larger subsamples and greater total sample volume. Polyethersulfone (0.8 μm) filters with glass fibre pre-filters (5 μm) were used and Longmire’s solution added for storage at ambient temperature for up to 1 week prior to DNA extraction. Therefore, successful filtration and greater detection in the present field study may be due to pond characteristics, sampling strategy, volume of water collected and filtered, filter material and pore size, inclusion of a pre-filter, use of a peristaltic pump, and/or preservation strategy.

Nonetheless, EP and filtration were inconsistent as standalone methods across all months and produced incongruent results to each other for individual months. Often, great crested newts were detected with one eDNA capture method but not the other across several months, or great crested newts were detected in one month but not the next month with a single eDNA capture method (Figure 3). Excluding control ponds (Ponds 1-4 and 24), each eDNA capture method produced different detection results to the other method within a given month for 17 ponds. Filtration produced incongruent results in seven ponds across several months, including three ponds where EP detected great crested newts at the start or end of the study period. EP produced incongruent results in 10 ponds across several months, including three ponds where filtration detected great crested newts at the start or end of the study period. During a single in-season month, 11 ponds that were positive for great crested newt DNA with filtration tested negative with EP, representing 11 potential failures to protect great crested newts under current legislation due to filtration not being accepted by regulators in line with best practice for eDNA analysis (but see Discussion below). This contrasts with two ponds that tested positive with EP and negative with filtration in a single in-season month. During a single out-of-season month, six ponds that were positive with filtration tested negative with EP, and five ponds that were positive with EP tested negative with filtration. Inconsistent eDNA results using EP and filtration at the same ponds were also observed by Rees et al. (2024).

Similar to in-season versus out-of-season detection, inconsistent detection between eDNA capture methods introduces some uncertainty and risk of false negatives with either eDNA capture method. This could be mitigated by using both eDNA capture methods at ponds being surveyed, but this would inherently increase survey time and cost. A compromise may be to resurvey ponds that are negative with EP kits using filtration kits, but this may still be unpopular with industry due to time and cost of collecting and analysing another sample as well as potential misalignment in detection sensitivity between filtration and conventional survey. Filtration may be substantially more sensitive and produce a positive result that would require population count surveys consisting of 4-6 site visits between mid-March and mid-June (with at least two visits between mid-April and mid-May), but great crested newts may not be observed by surveyors if present at low density. Heightened detection sensitivity could have benefits for overall conservation of this protected species but raises questions around the value of conserving small populations, similar to the value of non-breeding populations (Rees et al., 2023).

The performance of filtration and advantages for field surveyors and laboratories indicate that regulators should consider acceptance of filtration methods for non-regulated application at local scales. However, persistent barriers to uptake may be the high variability in filters (e.g. material, diameter, pore size, surface area) available to buy from scientific suppliers (Rees et al., 2023) as well as the strategies to preserve captured DNA (e.g. freezing, silica beads, preservation buffers) and storage conditions (e.g. temperature, duration), which can all influence the quality and quantity of eDNA extracted from samples and subsequently species detection (Hinlo et al., 2017; Jeunen et al., 2019; Minamoto et al., 2015; Peixoto et al. 2020; Spens et al., 2017). Concerns have also been expressed over effort required for manual filtration using syringes, filter clogging, negative eDNA results obtained with EP in previous surveys being undermined, a mismatch in detection sensitivity between filtration and ecologists performing population surveys, value of monitoring small and/or non-breeding populations, and implications for downstream analysis including proficiency and degradation testing, e.g. which filter type(s) and extraction protocols should be used by laboratories participating in testing (Rees et al., 2023, 2024). In particular, increased time and physical effort required for manual filtration using syringes (Andreou et al., 2023) may reduce the number of ponds (or sites) that can be surveyed in a single day and subsequently increase survey costs. In the present study, filtration time was ∼20 minutes compared to ∼10 minutes for EP, but using a peristaltic pump which may not always be logistically or financially feasible (Buxton et al., 2018b). Syringe filtration assisted by a silicone gun may be a suitable compromise to remove the need for a pump and increase volume filtered but reduce surveyor time and discomfort (Andreou et al., 2023). A standardised volume of water (e.g. 500 mL) to process with filtration will also be necessary to ensure consistency and comparability for regulatory monitoring whilst maintaining the same level of detection sensitivity (Andreou et al., 2023; Buxton et al., 2018b), but this may require use of a pre-filter or multiple filters that would need to be combined for analysis, or filter with larger pore size (Bruce et al., 2021; Rees et al., 2023, 2024).

The aforementioned barriers to uptake could be tested systematically through further field validation trials and ring testing with different ecological surveyors and laboratories. More broadly though, changing methods could have implications for the national great crested newt eDNA data set. Changing protocols in long-term monitoring schemes, particularly statutory or regulatory monitoring, is challenging due to the need for consistency and comparability of resulting data and potential for introducing bias. Indeed, this is a challenge frequently encountered when end users decide to incorporate eDNA survey into a monitoring scheme that has traditionally used other methods. Calibration is needed to assess and remove variability, such as that undertaken for acceptance of eDNA survey for great crested newt (Biggs et al., 2014, 2015) and determining the European Union Water Framework Directive classification for the lake fish biological element using eDNA metabarcoding (Sellers et al., 2024) in the UK. One approach to calibration would be to compare filtration and EP with just 90 mL of water from a pooled sample, with samples being collected from a large set of ponds with different physicochemical properties. Unpublished results suggest that filtration still outperforms EP for great crested newt detection using this small volume (Rees et al., 2023). However, it is unknown whether eDNA captured from 90 mL of water is sufficient for detection of other biodiversity (L. R. Harper et al., 2018). A study on a riverine fish community found that the number of species detected increased and community composition became more similar as more water was filtered, ranging from 10 mL to 1 L (Sakata et al., 2021). Alternatively, modelling is needed to assess changes in detection sensitivity and error rates resulting from methodological change, as applied to bighead carp and Asian carp eDNA monitoring in the United States (Song et al., 2019). Similar considerations are required for any changes to laboratory processing of great crested newt eDNA samples (Rees et al., 2023).

Additional evidence and a cost-benefit analysis of different filtration approaches (e.g. manual, pump-driven, on-site, off-site) versus EP is needed to overcome some of the aforementioned barriers and to determine the optimal filtration method for regulated application at national scale (Rees et al., 2023, 2024). To generate that evidence, two approaches may be considered. First, experiments, similar to the present study, are carried out that encompass waterbodies of known great crested newt status throughout the UK which span the full spectrum of physicochemical variability seen in these systems. The results would provide definitive answers for nationwide application. Alternatively, regulatory acceptance of results from projects similar to this study would allow local/regional great crested newt eDNA surveys to be conducted using filtration. This would require these projects to survey 10-20 ponds that capture the spectrum of physicochemical variability seen in the locality or region. With either approach, the filtration conditions (e.g. water volume, on-site versus off-site, manual versus pump, pore size, material etc.) should be taken into account. It would be beneficial for any future validation experiments to incorporate filter types and preservatives that have already been validated in other local contexts (Buxton et al., 2018b; Rees et al., 2024; this study) as well as those in common use (Bruce et al., 2021), e.g. Sterivex cartridges (Merck Millipore) and eDNA Dual Filter Capsules (Sylphium Molecular Ecology) with ethanol, RNAlater or lysis agents. Allowing various filter types and preservatives to be used nationally would not be robust and likely increase the rate of false negatives relative to that of current EP kits (Biggs et al., 2014, 2015) unless a study comparing filter types and preservatives at national scale was also undertaken.

### 4.4. Population size

Links between population size and amount of eDNA present continue to be inconsistent in the eDNA literature, with some studies finding a positive relationship and others finding no relationship. In natural systems, biotic and abiotic factors can influence the release, persistence, and degradation of eDNA, and these effects may be exacerbated in ponds which exhibit greater physical and chemical heterogeneity (L. R. Harper et al. 2019). Thomsen et al. (2012) observed a positive relationship between great crested newt density and eDNA concentration under controlled and natural conditions. Although Biggs et al. (2015) also observed this relationship between eDNA score (number of positive qPCR replicates) and adult counts, they found that a high eDNA score did not necessarily correspond to high adult counts. Buxton et al. (2017) found that adult numbers influenced eDNA concentration during the breeding season, but number of larvae and their size were the main drivers of eDNA concentration outside the breeding season. In our study, adult population size in the breeding season did not influence any permutation of eDNA score tested. However, different ponds were surveyed with different methods, including bottle, torchlight, netting and refuge searches (Table 1). Newt counts via torchlight can be compromised by water turbidity, vegetation cover, or weather conditions which affect visibility, and temperature which affects newt activity. Counts via refuge searching are also less reliable than other survey methods as even if newts are present, they may not be using refuges (Langton et al., 2001). Furthermore, methods for counting adults may only produce a minimum estimate of population size (Griffiths & Inns, 1998), and high adult counts may not correspond to breeding adults who are more likely to shed DNA. These factors as well as environmental conditions (Biggs et al., 2015) may be why no relationship was found here.

### 4.5. Insights possible by extending sampling approaches for great crested newt eDNA monitoring

In mainland Europe, where great crested newt eDNA monitoring is not subject to the same regulations as the UK, out-of-season sampling and filtration are frequently used. In Denmark, citizen scientists were recruited to collect filters from waterbodies between May and September for qPCR analysis of 14 species, including great crested newt (Knudsen et al., 2023). In Norway, water samples were collected in May and July then filtered off-site for qPCR analysis to examine presence of amphibians, including great crested newt, and chytrid fungus (*B. dendrobatidis*) in acidic ponds (Strand et al., 2022). Also in Norway, filtered water samples were analysed with droplet digital PCR assays for great crested newt, smooth newt (*Lissotriton vulgaris*) and chytrid fungus (*B. dendrobatidis*) to examine infection and potential declines (Taugbøl et al., 2021), and optimise field methods (Taugbøl et al., 2025). In France, qPCR analysis of filters collected in spring was used to investigate great crested newt occupancy alongside other methods (Priol et al., 2023).

The literature grows when studies with a whole community focus using eDNA metabarcoding are included. In Denmark, eDNA metabarcoding of filters collected in July was compared to conventional surveys for amphibians including great crested newt (Svenningsen et al., 2022). In Switzerland, eDNA metabarcoding of filters was used to examine distribution of great crested newt in relation to the invasive Italian crested newt (*Triturus carnifex*) and potential for hybridisation (Dufresnes et al., 2019). Also in Swtizerland, water samples were collected and filtered off-site for eDNA metabarcoding to examine amphibian distribution in urban areas (Charvoz et al., 2021). In the Netherlands, filters collected in April were analysed with qPCR analysis for chytrid fungus (*B. salamandrivorans*) and eDNA metabarcoding for amphibians to investigate possible transmission and infection (Davison et al., 2025).

These studies demonstrate the broader applicability of eDNA analysis for understanding great crested newt ecology, including distribution, occupancy, species interactions and disease susceptibility. Some studies examined the false positive and false negative error rate associated with their methods, but sampling was typically not repeated to examine the influence of survey timing and filtration was not compared to EP. Therefore, the future investigations described in sections 4.2 and 4.3 are still needed to ground truth and calibrate out-of-season survey and filtration, but this investment will allow for greater insights from great crested newt eDNA survey in the UK and internationally.

## 5. Conclusions

We have provided strong evidence that great crested newt eDNA detection is comparable or higher with filtration compared to EP, echoing many studies demonstrating this for other species and supporting a small number evidencing this for the great crested newt. Therefore, we recommend that this approach be included as an accepted eDNA capture method for great crested newt eDNA surveys by UK regulators. This could allow larger volumes of water to be processed for robust and reliable estimates of great crested newt presence. Extending the great crested newt eDNA survey season to August could allow more potential great crested newt sites to be surveyed to assess species presence (but not absence for reasons discussed) and identification of ponds that provide important habitat for great crested newts outside of the breeding season. This will also remove the logistical challenges and costs associated with completing sampling within 11 weeks and laboratory analysis within 10 working days from sample receipt. Furthermore, great crested newt eDNA surveys could be more frequently carried out alongside ecological surveying and monitoring for other species, which are typically surveyed from April to September/October or year-round with conventional methods or eDNA surveys using filtration. This would enable infrastructure projects to develop more effective mitigation measures as well as reduce time required from surveyors and survey costs.

We have highlighted future investigations that should be conducted to optimise great crested newt eDNA surveys and implement the changes desired by developers planning projects, consultants working in the field, and technicians working in the laboratory. However, change will only be possible with regulatory acceptance and incorporation of new evidence into established frameworks. Actions needed to extend the great crested newt eDNA survey season and allow the use of filtration for great crested newt eDNA capture are: 1) further consultation with regulators (i.e. Nature England, NatureScot, Natural Resources Wales), the Chartered Institute of Ecology and Environmental Management which oversees the ecological consultancy industry, amphibian conservation organisations, and eDNA service providers regarding the results of the present study and their interpretation, 2) consideration of the results from the present study together with other published evidence, 3) undertaking of a similar study to that reported here but including eDNA survey in February and March, and 4) validation experiments on filtration as a method of great crested newt eDNA capture at local, regional and national scale to enable regulatory acceptance at the corresponding spatial scale.

## Author contributions

Lynsey Harper and Bastian Egeter designed the study; Kat Stanhope managed the project on behalf of HS2 Ltd; Kirsten Harper performed the literature review; Suzie Platts, Laura Plant, Danielle Eccleshall, Luke Gorman and Kat Stanhope facilitated sample collection and provided supporting data; Rebecca Irwin and Michael Bennett performed the laboratory work; Ben Jones, Oliver Taylor and Lynsey Harper analysed the data; Lynsey Harper wrote the manuscript, which all authors revised and gave final approval for publication.

## Acknowledgements

We would like to thank HS2 Ltd for funding this work and permitting access to ponds. We are grateful to AtkinsRéalis employees for their efforts in the field to collect eDNA samples as well as conduct population size class and habitat suitability assessments. We are also grateful to Ben Parkinson for collating and providing data collected on behalf of HS2 Ltd. We appreciate constructive feedback received on the experimental design from Debbie Leatherland and Andy Nisbet (Natural England). We would like to thank end-users who responded to requests for grey literature, particularly Chris Troth (SureScreen Ltd) for sharing data.

## Conflict of interest

Lynsey Harper is employed by Natural England but they do not endorse this work. Some of this data has been published on the HS2 Learning Legacy website: https://learninglegacy.hs2.org.uk/document/environmental-dna-surveys-for-great-crested-newts-time-for-regulatory-changes/. Rebecca Irwin, Laura Plant, Andrew Briscoe and Bastian Egeter are employed by NatureMetrics Ltd, a for profit company dedicated to the analysis of environmental DNA. Suzie Platts is employed by Binnies, and Danielle Eccleshall and Luke Gorman are employed by AtkinsRéalis, for profit environmental consultancies. Kat Stanhope is employed by National Grid, a for profit electricity, natural gas and clean energy delivery company.

## Data availability

Scripts and corresponding data have been deposited in a dedicated Zenodo repository (https://doi.org/10.5281/zenodo.16544635).

## Notes

https://doi.org/10.5281/zenodo.16544635

### Summary of Updates

This version of the manuscript has been revised to address reviewers' comments for resubmission to a journal.

