## Supporting Information for "Extending sampling approaches for great crested newt (*Triturus cristatus*) eDNA monitoring"

### for

#### Supporting Methods

##### Literature review

From each source, the following information was extracted where possible: target species; environment(s) sampled; waterbody type; sample type; eDNA capture method; sample volume; sampling protocol; number of samples per waterbody; number of waterbodies sampled; number of sites surveyed; date of sampling; DNA extraction method; DNA amplification method; number of technical replicates performed per sample for DNA amplification; number of ponds testing positive for the target species; eDNA score; copy number (if applicable); direction and strength of correlation between eDNA score/copy number and any conventional population density estimates (if applicable); conventional population estimate; and any other metadata. For studies using ethanol precipitation (EP), the volume of water processed, percentage and volume of ethanol, molarity, pH and volume of sodium acetate (if applicable), storage temperature and duration, and centrifugation conditions were recorded. For studies using filtration, the volume of water processed, filter type (enclosed or open membrane), pore size, membrane material and diameter, preservation buffer (if applicable), storage temperature and duration were recorded. Information extracted from each source was summarised in Microsoft Excel (Appendix 1).

##### Occupancy modelling

The Griffin et al. (2020) model, implemented via the eDNAShinyApp package (Diana et al. 2021), enables false positive and false negative error rates to be estimated for eDNA surveys. This allows specification of covariates which may influence site occupancy and detection probability. We applied this model to our data, where the probabilities of occupancy ($\Psi$), Stage 1 (sample collection) true positive ($\theta_{11}$) and false positive ($\theta_{10}$) observations, and Stage 2 (laboratory analysis) true positive ($p_{11}$) and false positive ($p_{10}$) observations, were all considered with default prior settings from Griffin et al. (2020) of 0.9 for $\theta_{11}$ and $p_{11}$ and 0.1 for $\theta_{10}$ and $p_{10}$. However, different covariates cannot be specified at different levels with this model. Therefore, only $\theta$ was considered as a function of covariates in order to examine the effect of month and eDNA capture method on the probability of great crested newt eDNA being present in a sample. The package was run using 2,000 burn-in iterations, 10,000 iterations, 1 chain and 20 thinned iterations, with 'probability of site occupancy' set to 0.5, 'variance of probability of site occupancy' set to 4, 'variance of coefficients of probability of site occupancy' set to 0.25, and number of significant covariates' set to 2. Bayesian variable selection using an Add-Delete-Swap approach is automated within the eDNAShinyApp package using a Pólya-Gamma sampling scheme and the Markov Chain Monte Carlo (MCMC) algorithm presented in Griffin et al. (2020). Covariates were considered important if their respective posterior inclusion probability (PIP) value was greater than 0.5, indicating that they appear in more than 50% of model iterations and the 95% posterior credible intervals of their corresponding coefficients did not include zero.

The Dorazio and Erickson (2018) model, implemented with a Gibbs sampler via the msocc package for computational efficiency (Stratton et al. 2020), does allow different covariates to be specified at different levels, but only provides estimates of false negative error. Eight models were fit to our data, including a null model and models with different combinations of covariates (month and eDNA capture method) on $\theta$ and/or $p$. Each model was run with 1,000 burn-in MCMC samples and 10,000 MCMC samples. The widely applicable information criterion (WAIC; Watanabe & Opper, 2010) was derived for each model to compare the quality of models based on the overfitting–underfitting trade-off. Model-based estimates of θ in different months and with each eDNA capture method, and estimates of $p$ for each eDNA capture method were calculated from the intercepts and regression slopes of α and δ – the logit-scale parameters from which θ and $p$ are derived, respectively. These θ and $p$ estimates were calculated from all 10,000 (non-burn-in) MCMC samples of the joint posterior distribution, such that they could be summarised as medians and 95% credible intervals.

Neither the Griffin et al. (2020) or Dorazio and Erickson (2018) models account for repeat visits to the same site in the same way as a GLMM with a random effect does. Instead, each visit is treated as a unique site. Multi-method models, implemented via the RPresence package (MacKenzie & Hines, 2023), account for non-independence between sampling methods at shared, local scale sampling locations (Nichols et al., 2008). We estimated method-specific ($p_{EP}$ and $p_F$) and month-specific ($p_{April}$, $p_{May}$, $p_{June}$, $p_{July}$, $p_{August}$, $p_{September}$, $p_{October}$) detection parameters. We fixed occupancy (Ψ) and local occupancy (θ) to 1, because great crested newts were known to occur across all three sites and at all 20 ponds. It is not possible to implement the hierarchical dependence structure of eDNA surveys with this model and only false negative error rate is estimated.

#### Supporting Results

##### Literature review

All studies collected one water sample per pond comprising subsamples, but not all followed the established field protocol (Biggs et al., 2014) for great crested newt eDNA survey. The number of ponds included in each study ranged from 8-18 ponds across 1-3 sites. This does not include an unpublished data set of 130 ponds provided by SureScreen Ltd where the number of sites was unknown and location information was not available for a number of ponds. The eDNA capture conditions for EP and filtration, the DNA extraction method, and the amplification method were consistent across studies. The number of technical replicates per sample was 8 or 12 replicates for qPCR, and the number of positive ponds ranged from 1-42 ponds. Copy numbers were either not reported or not available, and very little metadata was available (Appendix 1).

##### eDNA capture method and survey season

For great crested newt status, the effect of eDNA capture method did not depend on survey season (regardless of whether September and October were included) and vice versa as the interaction term was not significant (Table S1). When all out-of-season months were included, the individual effect of survey season was significant but the individual effect of eDNA capture method was not (Table S1). Samples were more likely to be positive for great crested newt in-season than they were out-of-season, but samples were equally as likely to be positive for great crested newt with EP or filtration (Figures S1ai, bi). When September and October results were excluded, the individual effect of survey season was not significant whereas the individual effect of eDNA capture method was (Table S1). Samples were more likely to be positive with filtration than they were with EP, but samples were equally as likely to be positive for great crested newt in-season as they were in July and August (Figures S1aii, bii).

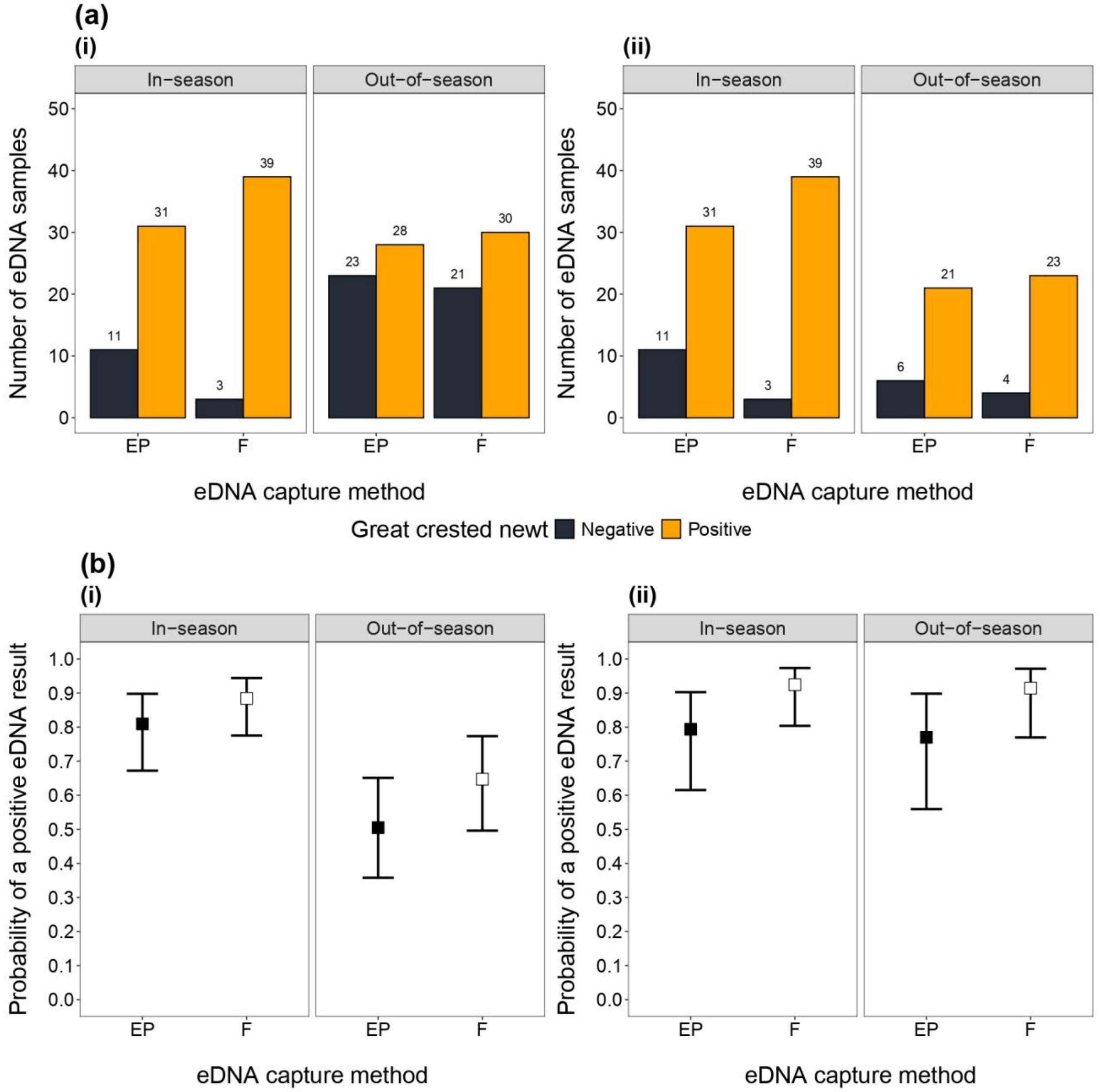


**Figure S1.** The effects of eDNA capture method and survey season on great crested newt detection. In **(a)**, bars show the number of samples that were positive (orange) and negative (black) for great crested newt in-season or out-of-season with EP and filtration (F) kits. The number of samples corresponding to each category are displayed above bars. In **(b)**, square points represent the model-predicted positive eDNA result probabilities with EP (black) and F (white) in-season or out-of-season, and error bars show the 95% confidence intervals for these predictions. Observed and predicted results are shown with September and October included **(i)** and excluded **(ii)** from out-of-season survey.

**Table S1.** Summary of GLMM and post hoc analysis performed to examine the influence of eDNA capture methods and great crested newt eDNA survey season (as individual and interacting effects) on the great crested newt status of samples, including and excluding the months of September and October. Significant p-values (<0.05) are in bold and df denotes degrees of freedom.

| September and October included? | GLMM | | | | | Post hoc analysis | | | | |
|---|---|---|---|---|---|---|---|---|---|---|
| | Formula | Variable | Effect | | | Levels | Mean probability ± SE | Pairwise comparison | | |
| | | | $X^2$ | df | P | | | Pair | z | P |
| Yes | great crested newt status ~ edna capture method + season + edna capture method : season | eDNA capture method : season | 3.271 | 1 | 0.071 | NA | NA | NA | NA | NA |
| | | eDNA capture method | 2.916 | 1 | 0.088 | EP | 0.675 ± 0.063 | EP : F | -1.692 | 0.091 |
| | | | | | | F | 0.789 ± 0.052 | | | |
| | | Season | 16.465 | 1 | **<0.001** | In-season | 0.851 ± 0.044 | In-season : Out-of-season | 3.820 | **<0.001** |
| | | | | | | Out-of-season | 0.578 ± 0.064 | | | |
| No | great crested newt status ~ edna capture method + season + edna capture method : season | eDNA capture method * season | 1.163 | 1 | 0.281 | NA | NA | NA | NA | NA |
| | | eDNA capture method | 5.618 | 1 | **0.018** | EP | 0.782 ± 0.069 | EP : F | -2.263 | **0.024** |
| | | | | | | F | 0.920 ± 0.039 | | | |
| | | Season | 0.081 | 1 | 0.777 | In-season | 0.873 ± 0.049 | In-season : Out-of-season | 0.285 | 0.776 |
| | | | | | | Out-of-season | 0.857 ± 0.059 | | | |

For eDNA score, the effect of eDNA capture method did not depend on survey season (regardless of whether September and October were included) and vice versa as the interaction term was not significant (Table S2). When all out-of-season months were included, the individual effects of survey season and eDNA capture method were both significant (Table S2). Filtration produced higher eDNA scores than EP, and eDNA scores were higher in-season than out-of-season (Figures S2ai, bi). When September and October results were excluded, the individual effect of survey season was not significant whereas the individual effect of eDNA capture method was (Table S2). Filtration produced higher eDNA scores than EP, but eDNA scores were comparable in July and August to in-season months (Figures S2aii, bii).

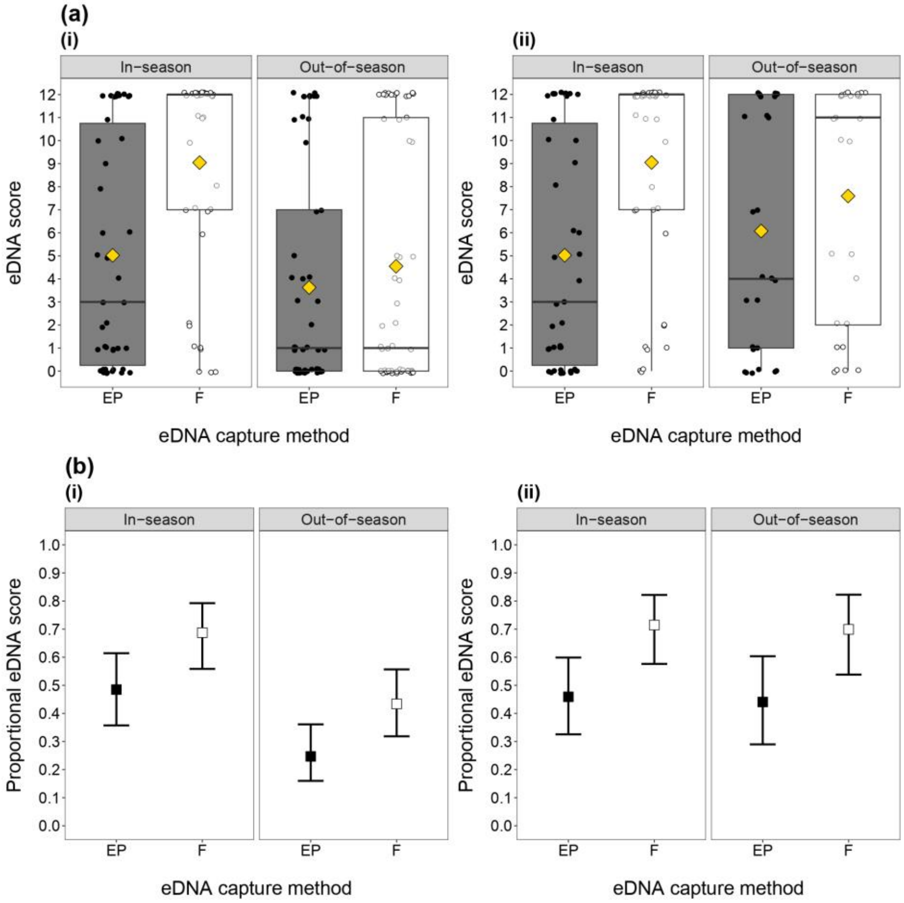


**Figure S2.** The effects of eDNA capture method and survey season on eDNA score. In **(a)**, the spread of eDNA scores with EP and filtration (F) kits in-season or out-of-season is represented by boxes which show the 25th, 50th and 75th percentiles, whiskers which show the 5th and 95th percentiles, and thick horizontal lines which show the median. Circular points (EP = black, F = white) represent the eDNA score for individual samples, and yellow diamonds represent the mean eDNA score for EP and F. In **(b)**, the square points represent the model-predicted proportional eDNA scores with EP (black) and F (white) in-season or out-of-season, and error bars show the 95% confidence intervals for these predictions. Observed and predicted results are shown with September and October included **(i)** and excluded **(ii)** from out-of-season survey.

**Table S2.** Summary of GLMM and post hoc analysis performed to examine the influence of eDNA capture methods and great crested newt eDNA survey season (as individual and interacting effects) on the proportional eDNA score of samples, including and excluding the months of September and October. Significant p-values (<0.05) are in bold and df denotes degrees of freedom.

| September and October included? | GLMM | | | | | Post hoc analysis | | | | |
|---|---|---|---|---|---|---|---|---|---|---|
| | Formula | Variable | Effect | | | Levels | Mean proportional eDNA score ± SE | Pairwise comparison | | |
| | | | $X^2$ | df | P | | | Pair | z | P |
| Yes | Proportional eDNA score ~ eDNA capture method + season + eDNA capture method : season | eDNA capture method : season | 3.068 | 1 | 0.080 | NA | NA | NA | NA | NA |
| | | eDNA capture method | 7.578 | 1 | **0.006** | EP | 0.357 ± 0.051 | EP : F | -2.716 | **0.007** |
| | | | | | | F | 0.565 ± 0.053 | | | |
| | | Season | 11.789 | 1 | **<0.001** | In-season | 0.590 ± 0.055 | In-season : Out-of-season | 3.371 | **<0.001** |
| | | | | | | Out-of-season | 0.334 ± 0.048 | | | |
| No | Proportional eDNA score ~ eDNA capture method + season + eDNA capture method : season | eDNA capture method : season | 1.625 | 1 | 0.203 | NA | NA | NA | NA | NA |
| | | eDNA capture method | 9.317 | 1 | **0.002** | EP | 0.450 ± 0.063 | EP : F | -2.974 | **0.003** |
| | | | | | | F | 0.707 ± 0.057 | | | |
| | | Season | 0.041 | 1 | 0.840 | In-season | 0.593 ± 0.058 | In-season : Out-of-season | 0.202 | 0.840 |
| | | | | | | Out-of-season | 0.575 ± 0.071 | | | |

#### Volume filtered

Volume filtered ranged from 125 mL to 2,500 mL across ponds (mean = 1,275 mL, median = 1,125 mL). There was no significant influence of volume filtered on eDNA score for samples collected with filtration kits (GLMM: $X^2_1 = 1.811$, $P = 0.178$). Samples with low volume filtered were capable of producing high eDNA scores and samples with high volume filtered also produced low eDNA scores (Figure S3).

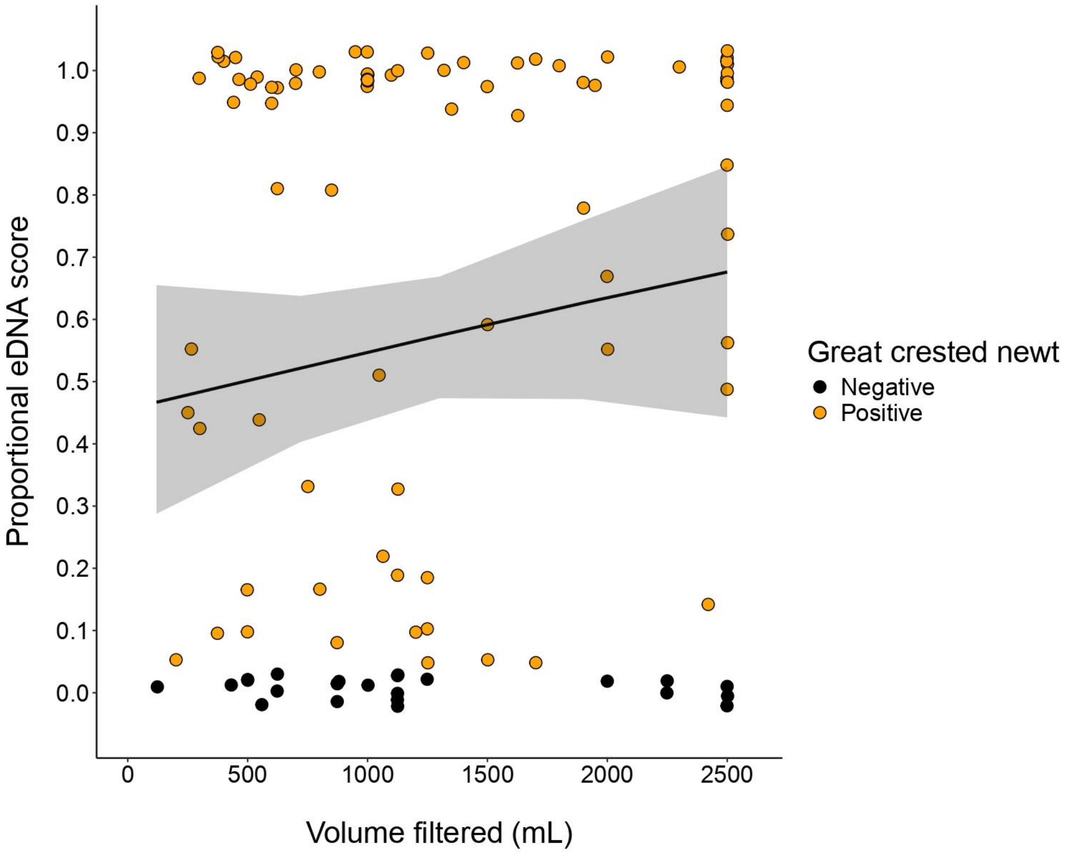


**Figure S3.** Scatter plot showing the volume filtered for each sample and corresponding eDNA score. Points are coloured by whether a sample was positive (orange) or negative (black) for great crested newt using filtration. Points are jittered to facilitate visualisation. The line represents the model-predicted proportional eDNA scores for different volumes, and the shaded area represents the 95% confidence intervals for these predictions.

#### Population size

There was no influence of peak adult counts or population class on corresponding eDNA score, average eDNA score, or peak eDNA score with EP or filtration (Table S3, Figure S4).

**Table S3.** Summary of GLMM analysis performed to examine the influence of peak adult counts and population class on the corresponding eDNA score for May, average eDNA score across in-season months, and peak eDNA score across in-season months for each pond. Significant p-values (<0.05) are in bold and df denotes degrees of freedom. Note that the GLMM for corresponding eDNA score with filtration experienced problems with underdispersion and convergence thus estimates could not be obtained.

| eDNA capture method | GLMM | | | | | Post-hoc analysis | | | | |
|---|---|---|---|---|---|---|---|---|---|---|
| | Formula | Variable | $X^2$ | df | P | Levels | Mean eDNA score ± SE | Pairwise comparison | | |
| | | | | | | | | Pair | z | P |
| Ethanol precipitation | Corresponding eDNA score ~ peak counts + population class | Peak count | 0.745 | 1 | 0.388 | NA | NA | NA | NA | NA |
| | | Population class | 2.676 | 2 | 0.262 | NA | NA | NA | NA | NA |
| | Average eDNA score ~ peak counts + population class | Peak count | 0.332 | 1 | 0.565 | NA | NA | NA | NA | NA |
| | | Population class | 1.628 | 2 | 0.443 | NA | NA | NA | NA | NA |
| | Peak eDNA score ~ peak counts + population class | Peak count | 0.144 | 1 | 0.704 | NA | NA | NA | NA | NA |
| | | Population class | 2.146 | 2 | 0.342 | NA | NA | NA | NA | NA |
| Filtration | Corresponding eDNA score ~ peak counts + population class | Peak count | NA | NA | NA | NA | NA | NA | NA | NA |
| | | Population class | NA | NA | NA | NA | NA | NA | NA | NA |
| | Average eDNA score ~ peak counts + population class | Peak count | 0.194 | 1 | 0.659 | NA | NA | NA | NA | NA |
| | | Population class | 0.158 | 2 | 0.924 | NA | NA | NA | NA | NA |
| | Peak eDNA score ~ peak counts + | Peak count | $-1.235 \times 10^{-10}$ | 1 | 1.000 | NA | NA | NA | NA | NA |

| eDNA capture method | GLMM | | | | | Post-hoc analysis | | | | |
|---|---|---|---|---|---|---|---|---|---|---|
| | Formula | Variable | $X^2$ | df | P | Levels | Mean eDNA score ± SE | Pairwise comparison | | |
| | | | | | | | | Pair | z | P |
| | population class | Population class | 3.138 x $10^{-11}$ | 2 | 1.000 | NA | NA | NA | NA | NA |

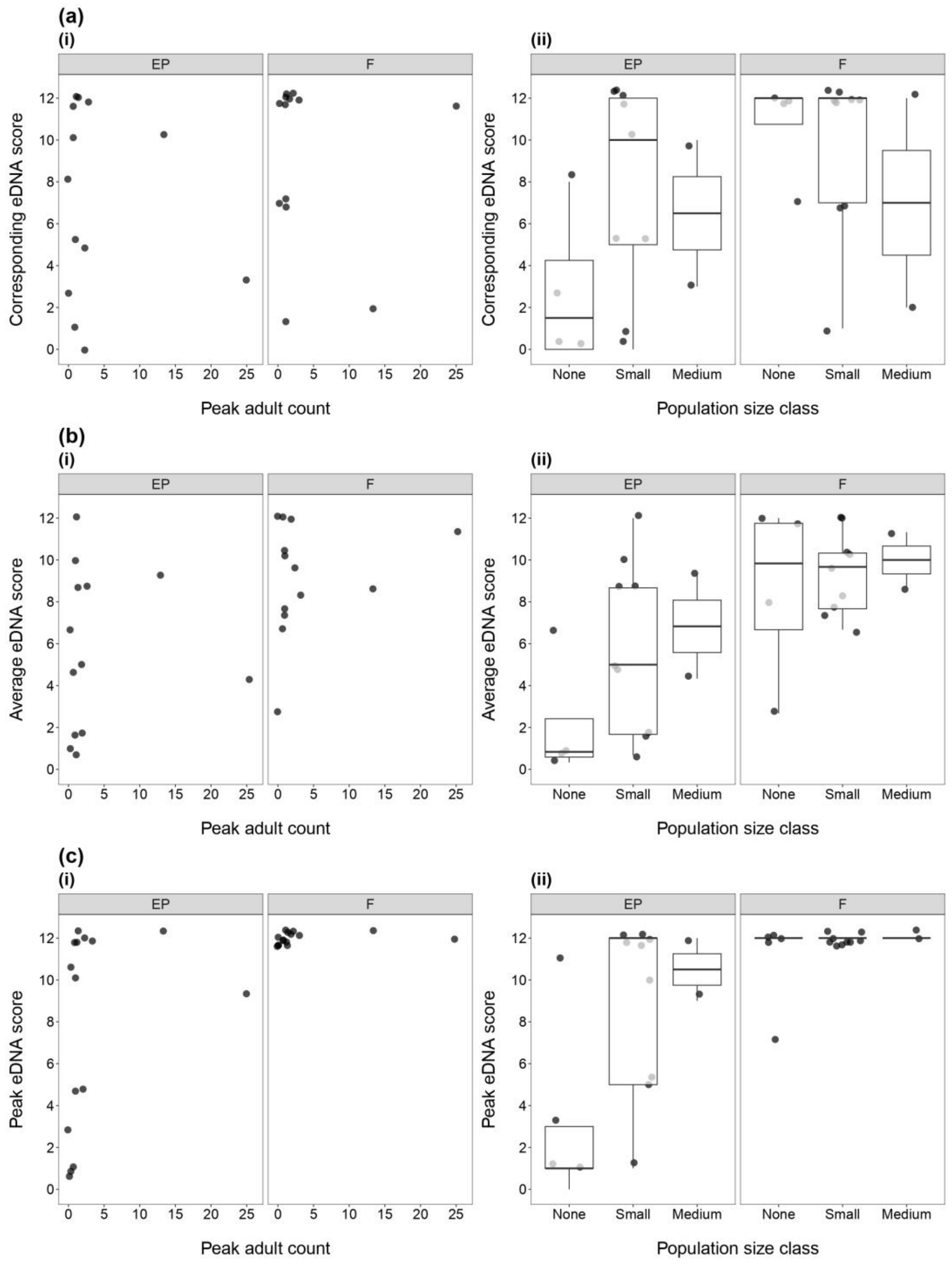


**Figure S4.** Plots showing the population size for each pond in relation to the **(a)** corresponding eDNA score for May, **(b)** average eDNA score across in-season months, or **(c)** peak eDNA score with ethanol precipitation (EP) or filtration (F) across in-season months. Population size is displayed as a scatter plot for peak adult count **(i)** or a box plot for population size class **(ii)**. Each point is a pond, boxes show the 25th, 50th and 75th percentiles, whiskers show the 5th and 95th percentiles, and thick horizontal lines show the median. Points are jittered to facilitate visualisation.

#### Appendices

Appendix 1. Summary of information extracted for each source identified by the literature review.

Appendix 2. Field study data set.
